# Genome-wide association analysis of dementia and its clinical endophenotypes reveal novel loci associated with Alzheimer’s disease and three causality networks of AD: the GR@ACE project

**DOI:** 10.1101/528901

**Authors:** Sonia Moreno-Grau, Itziar de Rojas, Isabel Hernández, Inés Quintela, Laura Montrreal, Montserrat Alegret, Begoña Hernández-Olasagarre, Laura Madrid, Antonio González-Perez, Olalla Maroña, Maitée Rosende-Roca, Ana Mauleón, Liliana Vargas, Asunción Lafuente, Carla Abdelnour, Octavio Rodríguez-Gómez, Silvia Gil, Miguel Ángel Santos-Santos, Ana Espinosa, Gemma Ortega, Ángela Sanabria, Alba Pérez-Cordón, Susana Ruiz, Nuria Aguilera, Juan Antonio Pineda, Juan Macías, Emilio Alarcón, Oscar Sotolongo-Grau, GR@ACE/DEGESCO consortium, Alzheimer’s Disease Neuroimaging Initiative, Marta Marquié, Gemma Montè-Rubio, Sergi Valero, Jordi Clarimón, Maria Jesus Bullido, Guillermo García-Ribas, Pau Pástor, Pascual Sánchez-Juan, Victoria Álvarez, Gerard Piñol-Ripoll, Jose Maria García-Alberca, José Luis Royo, Emilio Franco, Pablo Mir, Miguel Calero, Miguel Medina, Alberto Rábano, Jesús Ávila, Carmen Antúnez, Luis Miguel Real, Adelina Orellana, Ángel Carracedo, María Eugenia Sáez, Lluís Tárraga, Mercé Boada, Agustín Ruiz

**Affiliations:** Research Center and Memory clinic Fundació ACE. Institut Català de Neurociències Aplicades. Universitat Internacional de Catalunya. Barcelona. Spain; CIBERNED, Network Center for Biomedical Research in Neurodegenerative Diseases, National Institute of Health Carlos III, Spain; Grupo de Medicina Xenómica, Centro Nacional de Genotipado (CEGEN-PRB3-ISCIII). Universidade de Santiago de Compostela, Santiago de Compostela, Spain; CAEBI. Centro Andaluz de Estudios Bioinformáticos., Sevilla, Spain; Unidad Clínica de Enfermedades Infecciosas y Microbiología. Hospital Universitario de Valme, Sevilla, Spain; Dep. of Surgery, Biochemistry and Molecular Biology, School of Medicine. University of Málaga. Málaga. Spain; Memory Unit, Neurology Department and Sant Pau Biomedical Research Institute, Hospital de la Santa Creu i Sant Pau, Universitat Autònoma de Barcelona, Barcelona, Spain; Centro de Biologia Molecular Severo Ochoa (C.S.I.C.-U.A.M.), Universidad Autonoma de Madrid, Madrid, Spain; Instituto de Investigacion Sanitaria “Hospital la Paz” (IdlPaz), Madrid, Spain; Hospital Universitario Ramón y Cajal; Fundació per la Recerca Biomèdica i Social Mútua Terrassa, and Memory Disorders Unit, Department of Neurology, Hospital Universitari Mutua de Terrassa, University of Barcelona School of Medicine, Terrassa, Barcelona, Spain; Neurology Service ‘Marqués de Valdecilla’ University Hospital (University of Cantabria and IDIVAL), Santander, Spain; Laboratorio de Genética Hospital Universitario Central de Asturias, Oviedo; Instituto de Investigación Biosanitaria del Principado de Asturias (ISPA); Unitat Trastorns Cognitius, Hospital Universitari Santa Maria de Lleida, Institut de Recerca Biomédica de Lleida (IRBLLeida), Lleida, España; Alzheimer Research Center & Memory Clinic. Andalusian Institute for Neuroscience. Málaga. Spain; Unidad de Demencias, Servicio de Neurología y Neurofisiología. Instituto de Biomedicina de Sevilla (IBiS), Hospital Universitario Virgen del Rocío/CSIC/Universidad de Sevilla, Seville, Spain; Unidad de Trastornos del Movimiento, Servicio de Neurología y Neurofisiología. Instituto de Biomedicina de Sevilla (IBiS), Hospital Universitario Virgen del Rocío/CSIC/Universidad de Sevilla, Seville, Spain; CIEN Foundation, Queen Sofia Foundation Alzheimer Center, Madrid, Spain; Instituto de Salud Carlos III (ISCIII), Madrid, Spain; BT-CIEN; Department of Molecular Neuropathology, Centro de Biología Molecular “Severo Ochoa” (CBMSO), Consejo Superior de Investigaciones Científicas (CSIC)/Universidad Autónoma de Madrid (UAM); Unidad de Demencias. Hospital Clínico Universitario Virgen de la Arrixaca; Fundación Pública Galega de Medicina Xenómica-CIBERER-IDIS, Santiago de Compostela, Spain; CIBERNED, Center for Networked Biomedical Research on Neurodegenerative Diseases, National Institute of Health Carlos III, Ministry of Economy and Competitiveness, Spain; Centro de Investigación Biomédica en Red de Diabetes y Enfermedades Metabólicas Asociadas, CIBERDEM, Spain, Hospital Clínico San Carlos, Madrid, Spain; Research Center and Memory clinic Fundació ACE. Institut Català de Neurociències Aplicades. Universitat Internacional de Catalunya. Barcelona; Department of Neurology, Hospital Universitario Son Espases, Palma, Spain; Servei de Neurologia, Hospital Universitari i Politècnic La Fe; Centro de BiologiaMolecular Severo Ochoa (C.S.I.C.-U.A.M.), Universidad Autonoma de Madrid, Madrid, Spain; Instituto de Investigacion Sanitaria ‘Hospital la Paz’ (IdlPaz), Madrid, Spain; Hospital Universitario Ramón y Cajal; Madrid, Spain; BIOMICs, País Vasco; Centro de Investigación Lascaray. Universidad del País Vasco UPV/EHU; Neurology Service, Hospital Universitario La Paz (UAM), Madrid, Spain; Neurology Service, Marqués de Valdecilla University Hospital (University of Cantabria and IDIVAL), Santander, Spain; Hospital Donostia de San Sebastían; Fundación para la Formación e Investigación Sanitarias de la Región de Murcia; Servicio de Neurología-Hospital de Cabueñes-Gijón; Centro de Investigación y Terapias Avanzadas. Fundación CITA-alzheimer; Navarrabiomed; Servicio de Neurología-Hospital Universitario Central de Asturias, Oviedo; Barcelona βeta Brain Research Center – Fundació Pasqual Maragall; Unitat de Genètica Molecular. Institut de Biomedicina de València-CSIC; Unidad Mixta de Neurologia Genètica. Instituto de Investigación Sanitaria La Fe; Hospital Clínic Barcelona

**Author notes:** **Corresponding author**: Agustín Ruiz M.D. Ph.D., **Address**: Research Center. Fundació ACE. Institut Català de Neurociències Aplicades.C/ Marquès de Sentmenat, 57. 08029 Barcelona, Spain, **Email id**. **Alzheimer’s Disease Neuroimaging Initiative**: Data used in preparing this article were obtained from the Alzheimer’s Disease Neuroimaging Initiative (ADNI) database (adni.loni.usc.edu). As such, the investigators within the ADNI contributed to the design and implementation of ADNI and/or provided data but did not participate in the analysis or writing of this report. A complete listing of ADNI investigators can be found at http://adni.loni.usc.edu/wp-content/uploads/how_to_apply/ADNI_Acknowledgement_List.pdf.

**Keywords:** Alzheimer’s disease, vascular pathology, cerebral amyloid angiopathy, GWAS, biological pathway

## Abstract

**Background:** Genetics plays a major role in Alzheimer’s Disease (AD). To date, 40 genes associated with AD have been identified, although most remain undiscovered. Clinical, neuropathological and genetic variability might impact genetic discoveries and complicate dissection of the biological pathways underlying AD.

**Methods:** GR@ACE is a genome-wide study of dementia and its clinical endophenotypes that encompasses 4,120 cases and 3,289 controls from Spain. GR@ACE phenotypes were defined according to AD’s clinical certainty and the presence of vascular co-morbidity. To explore whether clinical endophenotypes reflect variation in underlying biological pathways, we first assessed the impact of known AD loci across endophenotypes to generate three loci categories. Next, we incorporated gene co-expression data and conducted pathway analysis on each category. To assess the impact of heterogeneity in the GWAS findings, the GR@ACE series were meta-analyzed with: 1) genotype-level data from dbGaP (N=21,235); and 2) summary statistics from IGAP Stages I and II (n=61,571 and n=81,455 respectively).

**Findings:** We classified known AD loci in three categories, which might reflect the disease clinical heterogeneity, from vascular and mixed forms to pure AD pathology. Immune system pathways were detected in all categories. Intriguingly, vascular processes were only detected as a causal mechanism in probable AD. A meta-analysis of GR@ACE with additional GWAS datasets revealed the *ANKRD31-rs4704171* signal in the *HMGCR* genomic region. We confirmed NDUFAF6-rs10098778 and *SCIMP*-rs7225151, which were previously detected by IGAP, to be suggestive signals. We also confirmed CD33-rs3865444 to be genome-wide significant.

**Interpretation:** The regulation of vasculature is a prominent causal component of probable AD. In that context, cerebral amyloid angiopathy, the unique identified link between the vascular and amyloid hypotheses, deserves further investigation. The GR@ACE meta-analysis revealed novel AD genetic signals. GWAS results are strongly driven by the presence of clinical heterogeneity in the AD series.

**Funding:** Grifols SA, Fundación bancaria “La Caixa”, Fundació ACE and ISCIII (Instituto de Salud Carlos III).

## Introduction

Dementia is an age-related clinical syndrome that devastates cognitive abilities and interferes in elderly people’s daily activities. Although its incidence is decreasing due to improvements to public health systems and control of cardiovascular risk factors,^1^ its prevalence is steadily increasing due to rising life expectancy of human populations.

Dementia is linked to many underlying pathologies, with Alzheimer’s Disease (AD) as the most common condition. AD brain autopsies commonly reveal three neuropathological hallmarks: amyloidosis, neurofibrillary tangles (NFTs), and cerebral amyloid angiopathy (CAA)^3^. However, the co-existence of other brain pathologies, especially cerebral vessel pathology, is the rule, and the number of “pure” AD cases is relatively small^4^. In that context, it has been proposed that there is a spectrum of this disease composed of a gradient of vascular and neurodegenerative features.^5^

Genetic factors play a pivotal role in AD etiology. In fact, two forms of the disease can be differentiated according to individual genetic background. The mendelian form is an uncommon disorder that mainly affects families with Early-onset AD (EOAD) (<65 years old), whereas the polygenic form is a complex disorder that mainly appears in sporadic cases with late-onset AD (LOAD) (>65 years old). Highly penetrant mutations detected in EOAD families have been pinpointed to three genes: *APP*,^6^ *PSEN1*,^7^ and *PSEN2*.^8^ The connection between earlier genetic and neuropathological findings promoted the establishment of the amyloid hypothesis as a potential causal mechanism for the disease.

LOAD heritability falls in the range of 25%–80%.^9,10^ *APOE* ε4 was the first to be discovered and still remains the strongest genetic risk factor for AD.^11^ The identification of additional genetic factors has only been feasible with the emergence of genome-wide association studies (GWAS) and large sequencing projects. Almost 40 genetic variants have been identified in these ways.^12,13,14^ Despite these advances, current genetic findings account for 31% of LOAD heritability.^15^ The missing heritability may be explained by several reasons: first, LOAD presents a complex genetic architecture; second, a lack of statistical power to detect uncharacterized variants with small effects; and third, the presence of ethnic differences together with clinical and neuropathological heterogeneity between AD cases.

The clinical and neuropathological variability of AD cases, comprising those with concomitant vascular brain disease to those with a pure AD phenotype, might be hampering the identification of functional categories of genes underlying differential biological routes to dementia. To gain insight on the causality networks behind AD clinical heterogeneity, we conducted the Genome Research at Fundacio ACE (GR@ACE) study, a GWAS of dementia and its clinical endophenotypes defined according to AD’s clinical certainty and the presence of vascular co-morbidity. GR@ACE is a unique genomic resource comprising a GWAS of the largest number of dementia cases diagnosed in a single memory clinic reported to date. First, we determined whether we could identify categories of known genes underlying clinical endophenotypes. Next, we explored whether these categories underpinned different biological routes. Finally to assess the impact of heterogeneity in GWAS findings and to address the need for more powerful and comprehensive AD genetic studies, we meta-analyzed the GR@ACE data with independent GWAS series.

## Methods

### Gr@ACE cohort and phenotype definitions

The GR@ACE study comprises 4,120 AD cases and 3,289 control individuals (table 1). Cases were recruited from Fundació ACE, Institut Català de Neurociències Aplicades (Catalonia, Spain). Diagnoses were established by a multidisciplinary working-group, including neurologists, neuropsychologists and social workers, according to the DSM-IV criteria for dementia and to the National Institute on Aging and Alzheimer’s Association’s (NIA-AA) 2011 guidelines for defining AD. In the present study, we considered AD cases, dementia individuals diagnosed with probable or possible AD at any moment of their clinical course.

**Table1.** GR@ACE demographic characteristic and endophenotype definitions.

| **Phenotype** | **Primary Diagnostic** | **Secondary Diagnostic** | **N** | **Mean Age ± (SD)** | **Women %** | **APOE ε4 %** |
| --- | --- | --- | --- | --- | --- | --- |
| Controls | -- | -- | 3289 | 54.3 ± 14.4 | 48.9 | 21.4 |
| VaD<sup>++</sup> | VaD | Pss AD | 373 | 80.1 ± 5.5 | 54.9 | 25.0 |
| VaD<sup>+</sup> | VaD/ Pss AD | VaD/ Pss AD | 1168 | 80.4 ± 6.3 | 65.0 | 32.8 |
| AD | Pr/Pss AD at any time in medical history | | 4120 | 79.0 ± 7.5 | 69.6 | 40.1 |
| AD<sup>+</sup> | Pr/Pss AD | Pr/Pss AD | 3797 | 79.2 ± 7.5 | 70.6 | 41.2 |
| AD<sup>++</sup> | Pr AD | Pr/Pss AD | 2611 | 78.8 ± 7.9 | 72.8 | 44.6 |
| AD<sup>+++</sup> | Pr AD | Pr AD | 1854 | 79.0 ± 8.0 | 74.6 | 47.0 |

VaD = Vascular Dementia; Pss AD = Possible AD; Pr AD = Probable AD.

We took advantage of this wide clinical definition to refine AD cases according to the degree of clinical certainty for AD phenotype and the presence of vascular comorbidity. This approach was feasible due to Fundació ACE’s endorsement of both a primary and a secondary etiologic diagnosis, as well as routine follow-up evaluations^16^ (see appendix). Using the entire clinical chart of each subject, we differentiated five clinical sub-groups of patients representing the GR@ACE endophenotypes: 1) the AD^+++^ endophenotype comprises individuals with a last clinical diagnosis of probable AD in both primary and secondary diagnoses (n = 1,854); 2) the AD^++^ includes individuals diagnosed with probable AD either in the primary or the secondary diagnosis (n = 2,611); 3) the AD^+^ encompasses patients diagnosed with probable or possible AD either in the primary diagnosis or in the secondary diagnosis (n = 3,797); 4) the VaD^+^ includes patients diagnosed with vascular dementia (VaD) or possible AD in the primary diagnosis (n = 1,168); and 5) the VaD^++^ comprises patients diagnosed with probable or possible vascular dementia in the primary diagnosis (n = 373) (table 1). The appendix includes a flow chart diagram detailing the endophenotype construction and a complete description of the neurological and neuropsychological assessments supporting the clinical diagnosis. VaD patients were defined according to NINDS-AIREN criteria.^17^

Control individuals were recruited from three centers: Fundació ACE (Barcelona, Spain), Valme University Hospital (Seville, Spain) and the Spanish National DNA Bank Carlos III (University of Salamanca, Spain) (www.bancoadn.org). Written informed consent was obtained from all participants. The Ethics and Scientific Committees have approved this research protocol (Acta 25/2016. Ethics Committee. H. Clinic i Provincial, Barcelona, Spain).

### GWAS genotyping, quality control, imputation and statistical analysis

Participants were genotyped using the Axiom 815K Spanish Biobank array (Thermo Fisher). Genotyping was performed in the Spanish National Center for Genotyping (CeGEN, Santiago de Compostela, Spain) (appendix).

We removed samples with genotype call rates below 97%, excess heterozygosity, duplicates, samples genetically related to other individuals in the cohort or sample mix-up (PIHAT > 0 1875). If a sex discrepancy was detected, the sample was removed unless the discrepancy was safely resolved. To detect population outliers of non-European ancestry (>6 SD from European population mean), principal component analysis (PCA) was conducted using SMARTPCA from EIGENSOFT 6.1.4 (figure 1) (appendix).

**Figure 1.**
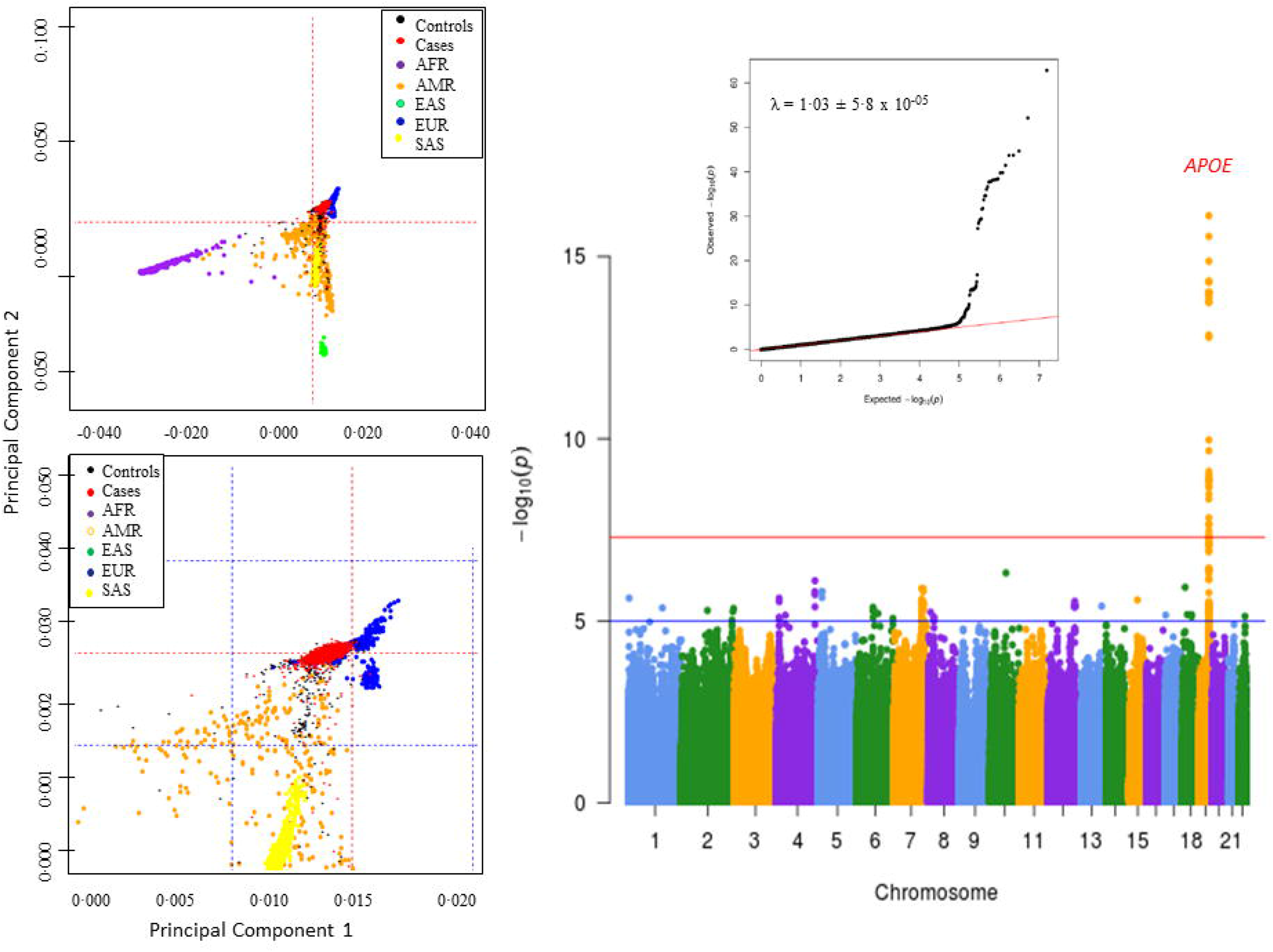
Results of genome-wide association analysis for the GR@ACE dataset (n = 7,409). Principal component analysis and QQplot.

We removed variants with a call rate <95% or that grossly deviated from Hardy-Weinberg equilibrium in controls (P-value ≤ 1x10^−4^), markers with different missing rate between case and control (P-value < 5 × 10^−4^ for the difference) or minor allele frequency (MAF) below 0·01. Imputation was carried out using Haplotype reference consortium (HRC) panel in Michigan Imputation servers (https://imputationserver.sph.umich.edu). Only common markers (MAF>0 01) with a high imputation quality (*R*^2^ >0·30) were selected to conduct downstream association analyses. Genome-wide association analyses were conducted for genotype dosages using an additive genetic model with PLINK 1.9. A model including the top four PCs as covariates was used for the discovery stage (see appendix).

### Genetic exploration of GR@ACE clinical endophenotypes and enrichment analysis

To explore whether differential gene categories were operating under GR@ACE clinical endophenotypes, we first evaluated the effect size change for known LOAD genetic variants (MAF>1%) on each phenotype. Effect size change represents the difference between variant odds ratio (OR) and null-effect (OR=1), and provides information about the strength of the association. To establish categories, we calculated the global effect change, defined as the effect change difference between the extreme endophenotypes (VaD^++^ vs AD^+++^). Thus, Category A includes variants with an increase in the association effect from VaD^++^ to AD^+++^ endophenotypes and a global effect change > 0 05; Category B, variants have an increase in the association effect from AD^+++^ to VaD^++^ and a global effect change > 0 05. Category C comprises variants not fulfilling criteria for categories A or B. Global effect changes are reported in the appendix. Finally, we assessed the biological pathways underlying each category. We incorporated data from gene co-expression using the GeneFriends tool (http://genefriends.org/) and performed pathway analysis using the overrepresentation enrichment method in WebGestalt (http://www.webgestalt.org/option.php). Additional sub-analysis of the genetic variants in Category C was performed. To validate previous gene classification per categories, which strongly determines the pathway analysis results, we conducted a stringent subanalysis. See the appendix for further description.

### Meta-analysis: datasets, association analysis and biological interpretation

To identify new loci associated with AD, we combined the GR@ACE GWAS dataset and its endophenotypes with: 1) genotype-level data from nine additional GWAS series (N = 21,235) available through dbGaP (https://www.ncbi.nlm.nih.gov/gap) that we processed by applying identical quality control and imputation procedures to those described for the GR@ACE cohort (appendix); 2) aggregated summary statistics publicly available from the International Genomics Alzheimer’s Project (IGAP) (http://web.pasteur-lille.fr/en/recherche/u744/igap/igap_download.php).^18^ See the appendix for further description of the meta-analysis cohorts. Meta-analyses were conducted using the inverse variant method in METAL software (https://genome.sph.umich.edu/wiki/METAL). The LD Score calculations, clumping and conditional analysis are described in appendix. Following, we conducted gene expression quantitative trait locus (eQTL) analysis to link meta-GWAS top signals to genes (see appendix).

### Results GR@ACE genome-wide association study

After quality control and imputation, the GR@ACE study encompassed 7,409 unrelated individuals from the Spanish population and 7.7 million variants (λ_GC_ = 1·03). The *APOE*-rs429358 marker was the only one to have a genome-wide significant (GWS) association [OR = 2.27 (2 06 – 2·50) p = 1·25 x 10^−62^] (figure 1). Four additional LOAD variants displayed statistically significant evidence of replication (BIN1-rs6733839, *MAPT*-rs2732703, *MS4A2*-rs983392, and *PICALM*-rs10792832) and nine additional markers presented a consistent direction for the effect (appendix). *MAPT* marker association was mainly driven by *APOE* ε4 non-carriers (appendix). Genome-wide results for GR@ACE endophenotypes are reported in appendix section and table 3.

### Genetic exploration of GR@ACE clinical endophenotypes and enrichment analysis

To explore whether GR@ACE clinical endophenotypes reflect variations in the underlying biological pathways driving dementia, we classified LOAD genetic variants in three categories. Category A comprised variants strongly related to the purest form of clinical AD (i.e., subjects with probable AD in primary and secondary diagnoses). The most prominent locus of this category was APOE-rs429358 [AD^+++^ OR = 2·92 (2·60 – 3·27), p-value = 9·26 x 10^−75^; VaD^++^ OR (95%) = 1·27 (1 02 – 1·59), p-value 0 04]. Other loci included in category A were *CR1, BIN1, MEF2C, MS4A2, PICALM, MAPT* and *CD33*. In contrast, category B comprised variants with the strongest effect observed in subjects with AD mixed with vascular disease (*SORL1, ADAM10, CASS4, ATP5H*, and *ACE*) (appendix). Category C comprised a group of variants with effects in all clinical endophenotypes. Figure 2 shows the enrichment trend per marker and by category.

**Figure 2.**
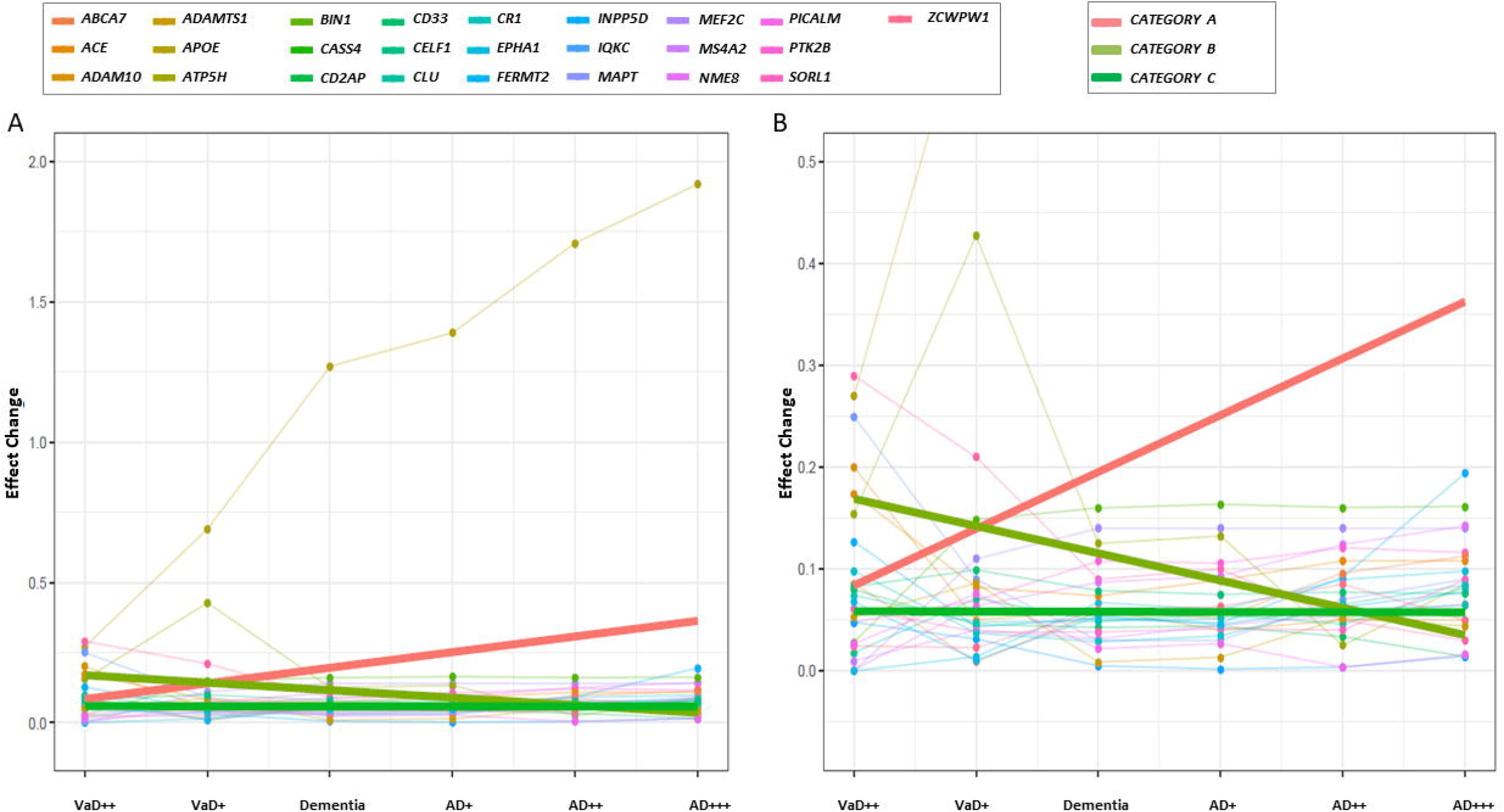
A) Enrichment trend per genetic marker and gene category across GR@ACE endophenotypes. B) Graph centered in effect change range 0 – 0.5. Enrichment trend per category was obtained applying a linear regression using ggplot2 in R.

Next, we explored biological pathways for each gene category. Note that the regulation of vasculature development and blood vessel morphogenesis were only detected for genes in category A, which is more closely related to pure AD (p =2 03 x 10^−7^, p = 1·90 x 10^−6^, respectively) (table 2). Additional categories indicated immune system pathways (Category B, p = 2 07 x 10^−7^; Category C, p = 5·77 x 10^−15^) (table 2). Finally, with the aim of validating previous results, we conducted a sub-analysis by classifying LOAD genetic variants with more stringent classification criteria (widely described in appendix). Again, *APOE, CR1, MEF2C, MS4A2* and *PICALM* loci were found in category A; *SORL1* and *CASS4* were in category B; and additional AD loci were in category C. The appendix presents the linear effect trends per variant. Regulation of vasculature development was exclusively identified as the top pathway in Category A (p = 214 x 10^−7^), when we restricted the analysis to include those loci co-expressing with, at least, 4 LOAD genes (appendix). Sub-analysis for Category C is shown in appendix.

**Table 2.** Top ten biological pathways per gene category.

| Gene Ontology Pathway | Top 10 co-regulated pathways for Category A | P-value |
| --- | --- | --- |
| GO:1901342 | regulation of vasculature development | $2.03 \times 10^{-7}$ |
| GO:0060326 | cell chemotaxis | $2.59 \times 10^{-7}$ |
| GO:0048771 | tissue remodeling | $6.77 \times 10^{-7}$ |
| GO:0050865 | regulation of cell activation | $1.14 \times 10^{-6}$ |
| GO:0007159 | leukocyte cell-cell adhesion | $1.21 \times 10^{-6}$ |
| GO:0048514 | blood vessel morphogenesis | $1.90 \times 10^{-6}$ |
| GO:0003012 | muscle system process | $2.54 \times 10^{-6}$ |
| GO:0002764 | immune response-regulating signaling pathway | $3.48 \times 10^{-6}$ |
| GO:0032103 | positive regulation of response to external stimulus | $3.91 \times 10^{-6}$ |
| GO:0010959 | regulation of metal ion transport | $4.36 \times 10^{-6}$ |
| Gene Ontology Pathway | Top 10 co-regulated pathways for Category B | P-value |
| GO:0009620 | response to fungus | $2.02 \times 10^{-7}$ |
| GO:0050886 | endocrine process | $3.58 \times 10^{-7}$ |
| GO:0002443 | leukocyte mediated immunity | $5.47 \times 10^{-7}$ |
| GO:0050865 | regulation of cell activation | $1.52 \times 10^{-5}$ |
| GO:0031349 | positive regulation of defense response | $8.42 \times 10^{-5}$ |
| GO:0032103 | positive regulation of response to external stimulus | $1.00 \times 10^{-4}$ |
| GO:0002250 | adaptive immune response | $1.30 \times 10^{-4}$ |
| GO:0098542 | defense response to other organism | $2.00 \times 10^{-4}$ |
| GO:1901568 | fatty acid derivative metabolic process | $2.24 \times 10^{-4}$ |
| GO:0050900 | leukocyte migration | $2.57 \times 10^{-4}$ |
| Gene Ontology Pathway | Top 10 co-regulated pathways for Category C | P-value |
| GO:0007159 | leukocyte cell-cell adhesion | $5.77 \times 10^{-15}$ |
| GO:0050865 | regulation of cell activation | $4.37 \times 10^{-14}$ |
| GO:0002764 | immune response-regulating signaling pathway | $1.33 \times 10^{-12}$ |
| GO:0002253 | activation of immune response | $3.96 \times 10^{-12}$ |
| GO:0002443 | leukocyte mediated immunity | $4.34 \times 10^{-12}$ |
| GO:0002274 | myeloid leukocyte activation | $7.78 \times 10^{-12}$ |
| GO:0002250 | adaptive immune response | $1.24 \times 10^{-11}$ |
| GO:0002263 | cell activation involved in immune response | $7.07 \times 10^{-11}$ |
| GO:0022407 | regulation of cell-cell adhesion | $5.40 \times 10^{-9}$ |
| GO:0070661 | leukocyte proliferation | $1.22 \times 10^{-8}$ |

### Meta-analysis of GR@ACE study with other datasets and eQTL analysis of Meta-GWAS signals

To look for new AD loci, we first combined the GR@ACE dataset with nine additional genomic databases that had genotypic level data available. Subtle genomic inflation was detected, mainly explained by polygenicity ((λ_GC_ = 1·10; LD score Intercept = 1·04). Five regions were associated with LOAD (figure 3); of these, four (APOE-rs429358, PICALM-rs10792832, MS4A2-rs983392 and BIN1-rs6733839) have been previously linked to AD, and one is a new GWAS significant finding [*ANKDR31*-rs4704171; OR = 1·19 (1·12 - 1·27); p = 2·78 x 10^−8^] (table 3). Forest plot for *ANKDR31*-rs4704171 is provided in appendix.

**Figure 3.**
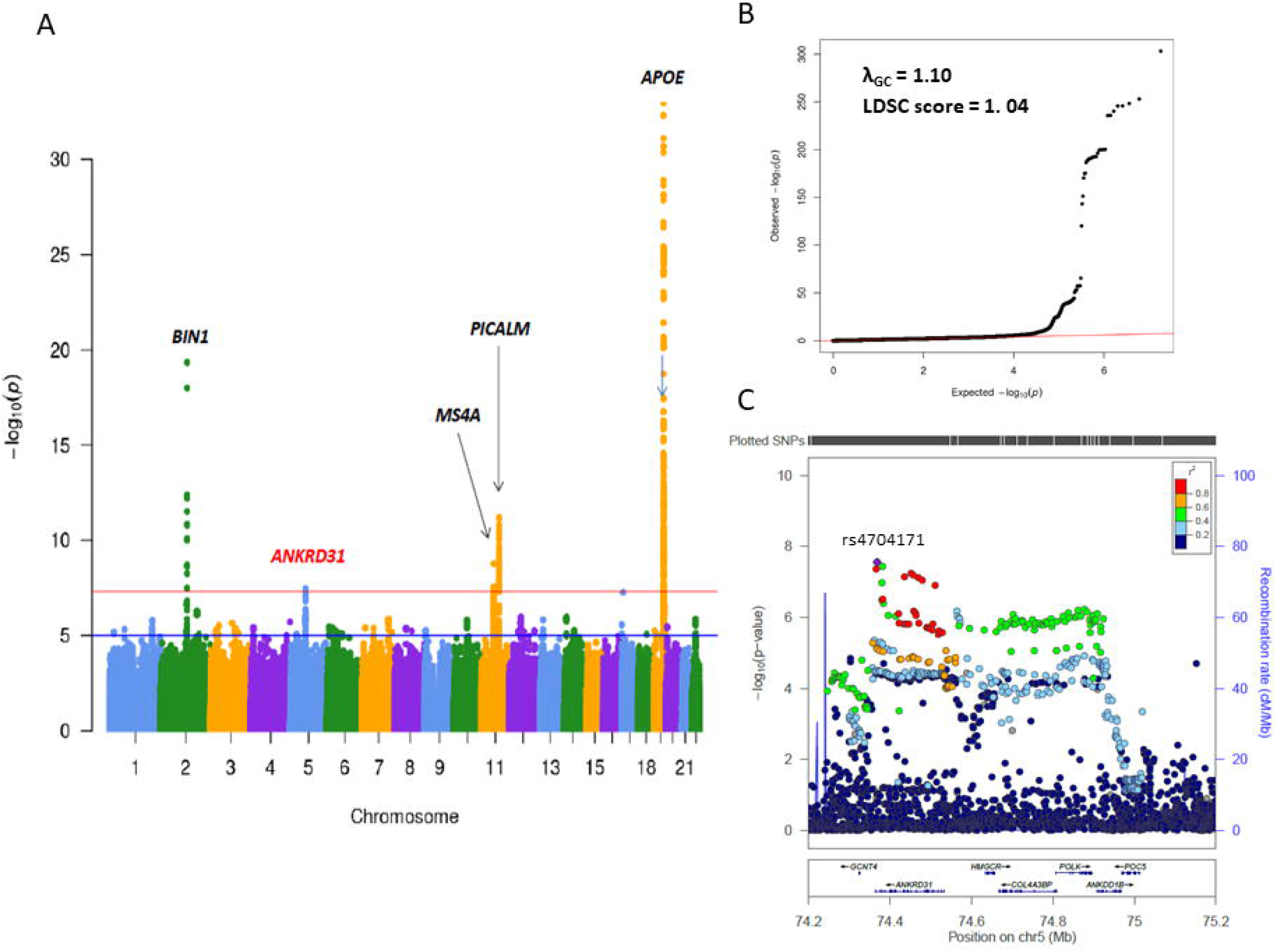
A) Results of genome-wide association analysis for GR@ACE meta-analysis with nine additional databases (n = 21,235). B) QQplot. C) Associations of the region centered on rs4704171 located in the *ANKRD31* locus and containing the *HMGCR* locus.

**Table 3.** Association results for lead single-nucleotide polymorphisms reaching genome-wide significance.

| Marker | Near Locus | Position<br>Chr:bp | Major/Minor<br>Allele | MAF | OR (CI95%) | P-value | Discovery stage |
| --- | --- | --- | --- | --- | --- | --- | --- |
| rs117834366 | *CNTNAP2* | 7:147634891 | G/A | 0.011 | 6.03<br>(3.22 – 11.2) | 1.91 x 10<sup>-8</sup> | GR@ACE VaD<sup>++</sup> |
| rs4704171* | *ANKRD31* | 5:74368254 | T/C | 0.123 | 1.19<br>(1.12 – 1.27) | 2.78 x 10<sup>-8</sup> | GR@ACE + dbGaP |
| rs10098778* | *TP53INP1/*<br>*NDUFAF6* | 8:95992020 | C/T | 0.470 | 1.07<br>(1.04 – 1.09) | 2.54 x 10<sup>-8</sup> | GR@ACE + IGAP<br>I&II |
| rs7225151* | *SCIMP* | 17:5137047 | G/A | 0.126 | 1.11<br>(1.07 – 1.15) | 1.12 x 10<sup>-8</sup> | GR@ACE<br>AD<sup>+++</sup> + IGAP I&II |

\*p-values were obtained using the fixed effect inverse-variant method. The threshold for genome-wide significance was 5 x 10<sup>-8</sup>. Position = GRCh37/hg19 coordinates; chr:bp = chromosome:base pair; MAF = Minor allele frequency obtained from the GR@ACE study.

Then, we conducted a genome-wide meta-analysis combining the GR@ACE study with IGAP stage I summary statistics (λ_GC_ = 1·09; LD Score Intercept = 1·03). We identified 13 LOAD genomic regions reaching GWS. Among these, CD33-rs3865444 which did not reach GWS in the IGAP meta-analysis, was significantly associated with LOAD [OR = 0·92 (0·89 – 0·95); p = 3·61 x 10^−8^]. We detected a suggestive signal in *HBEGF*-rs4150233 [OR = 0·92 (0·90 – 0·95); p = 5·10 x 10^−8^], previously identified by transethnic GWAS^19^ (see appendix).

Next, meta-analysis of the whole GR@ACE dataset with IGAP I and II summary statistics enabled the identification of NDUFAF6-rs10098778 [OR = 1 06 (1 04 - 1 09); p = 2·54 x 10^−8^). When we combine GR@ACE AD^+++^ endophenotype with IGAP I and II a GWS was detected in SCIMP-rs7225151 [OR = 1·11 (1·07 – 1·15); p = 1·12 x 10^−8^] (table 3) (appendix). Both signals have been previously reported as genome-wide suggestive signals by IGAP,^18^ and recently, *SCIMP* was significantly associated with AD.^14^

Finally, to identify candidate genes and potential causal variants within novel genome-wide regions, we conducted cis-eQTL mapping. Gene-mapping pointed to three genes with cortical expression and three additional ones expressed in blood. Further description is provided in appendix section.

## Discussion

We present a comprehensive and large genome-wide association study of AD dementia cases, including also its clinical endophenotypes. We showed differential biological classes underlying clinical endophenotypes and demonstrated that the composition of differential sub-groups of AD patients impacts GWAS discoveries. The GR@ACE study represents a unique genomic resource because all affected cases were diagnosed in a single memory clinic using the same screening and diagnostic techniques. This might limit potential sources of clinical variation between study participants, as has been recently demonstrated in a large meta-GWAS.^9^

Based on the increase in evidence suggesting that vascular brain pathology can act concomitantly with AD to produce more rapid cognitive decline,^20^ we explored the effect of known LOAD loci across different levels of vascular burden in dementia patients using only clinical definitions. Our basic idea was to dissect, from a molecular point of view, the model previously proposed by Viswanathan et al.^5^ We observed the existence of three categories of loci, which might reflect the disease’s clinical heterogeneity, from vascular and mixed forms to a more “pure” AD phenotype. Intriguingly, we detected vascular processes to be the main causal mechanism in clinically pure AD and we found the immune system pervasively detected across the three categories. Although both pathways have been previously associated with LOAD by network analysis,^21^ this is the first study to show that the association with the vascular system is conducted by AD-specific clinical subgroup. Despite these findings, replication in an independent and large single-site GWAS cohort might help contrast the proposed loci classification, which was based on the clinical endophenotypes of the GR@ACE cohort.

Silent changes occur in brain microvasculature during AD progression. In fact, CAA is a well-recognized AD pathological feature characterized by the accumulation of amyloid proteins, mainly Aβ_1-40_, in the walls of small cerebral vessels. CAA has been proposed to compromise the perivascular drainage of Aβ from the brain to the peripheral system.^22^ Almost all AD brains harbor CAA pathology to some extent, although *in vivo* most CAA cases remain undiagnosed, even when using the validated Boston criteria.^23^ Mendelian mutations of the *APP* gene have been found in both CAA and AD.^6,27^ *APOE* ε4 and *CR1* have been associated with an increased risk of CAA.^24,25^ In particular, distinct AD loci have been associated with capillary and non-capillary CAA.^26^ Between them, *APOE* ε4 was strongly related to capillary CAA^26^. These links make it conceivable a potential genetic overlap between CAA and AD, and suggest that CAA pathology could represents an underlying process for AD. In that context, we think that intrinsic alterations to the vasculature could contribute to disease pathogenesis in more pure forms of AD, explaining our results. Conversely, in AD individuals with evident cerebrovascular lesions comprising mixed forms, the additional role of cardiovascular risk factors, i.e., hypertension, atherosclerosis or arteriosclerosis, should be considered, as these could point to a systemic pathological state leading to vascular damage and dementia. This would accord with the limited genetic correlation between neurodegenerative and other neurologic disorders,^9^ as well as with results coming from heterochronic parabionts in aging models.^28^

Understanding the role of vasculature pathology in AD seems a pertinent step. In that scenario, CAA would be a key AD hallmark. CAA represents the unique identified link between the vascular and amyloid hypotheses, but it has been completely neglected in the original hypothesis formulation.

From a clinical point of view, placing each patient somewhere along the disease spectrum proposed by Viswanathan is complex.^5^ A deep understanding of heterogeneity in AD seems necessary to design better genetic studies, which must drive the discovery of novel loci and, ultimately, innovative targets for AD therapies. In this study, we explored how clinical heterogeneity might impact GWAS findings by integrating distinct GWAS datasets with either the GR@ACE cohort as a whole or its endophenotypes. We found several new GWS signals that seem strongly dependent on the sample composition. For example, after combining IGAP Stages I and II with the entire GR@ACE dataset, we identified genetic signals in the *NDUFAF6* genomic region. When this exercise was conducted using GR@ACE endophenotypes, the *NDUFAF6* signals disappeared. It is tempting to speculate that studying per endophenotypes has reduced statistical power. However, at this point, with a smaller sample size, the *SCIMP* signal was detected using the clinically “pure” AD GR@ACE endophenotype, suggesting that a purer AD dataset without clinical mixed dementia cases would be necessary to safely replicate this finding. We think that using the specific clinical subgroups of the AD population empowered this study to detect genes associated with specific disease axes.

An alternative strategy is taking advantage of clinical heterogeneity. Specifically, heterogeneity might play a dual role in genetic studies. Although it might decrease the power to detect genes associated with more specific clinical subgroups, incorporating detailed clinical AD definitions can also promote identifying genes shared with other conditions or co-pathologies such as SVD. In fact, this was the case for the *ATP5H* loci, which was previously found to be associated with AD^29^ and more recently found in relation to SVD.^30^ We think that the same applies to the *ANKRD31* findings. *ANKRD31* encodes a protein containing ankyrin-repeats, and has been involved in neurodevelopmental disorders ^31^. Of note, the brain eQTL of a linc-RNA, located 1.6kb from the *HMGCR* locus and residing in the *COL4A3BP* gene, was mapped for *ANKRD31* GWAS signals. The *HMGCR* locus is one of the most important co-regulators of cholesterol biosynthesis, and it is a therapeutic target of statins. The *COL4A3BP* gene is involved in lipid transport.^32^ Several studies have linked *HMGCR* polymorphisms and AD risk or age at onset for AD,^33^ and cholesterol pathway has been identified such as biological route shared between AD and small vessel disease. Interestingly, markers in the *POLK* locus, located in the same disequilibrium block of *ANKDR31* (figure 3), jointly conferred risk for AD and plasma levels of LDL.^34^ Taken together, these findings support the role of this genomic region in AD. The reported genetic signal should be considered a highly probable finding, although independent replications are still required.

*NDUFAF6* and *SCIMP* signals have presented suggestive evidence of association in IGAP studies.^12,18^, and recently, *SCIMP* reached GWS^14^. In the present work, *NDUFAF6* genomic region reached GWS for the first time, and we detected that *SCIMP* signal was mainly conducted by specific group of AD cases. The *NDUFAF6* genomic region, containing the *TP53INP1* locus, was first associated with AD in a gene-based analysis,^35^ and it has been involved in mitochondrial function. The *SCIMP* genomic region influences several eQTLs, from uncharacterized cortical lncRNA to blood eQTLs in *SCIMP* or its neighbor, the *RABEP1* locus, both of which are associated with immune system function.^36,37^ The *CD33* locus remains a controversial LOAD locus due to large meta-GWAS were unable to replicate previous this signal,^18^ but here it reached GWS. We previously proposed that the cryptic population sub-structure could explain the divergent observations for this locus.^38^

Note that the lack of definitive neuropathological data for AD cases used in this project is a severe limitation of the present study. Clinical definitions have important uncertainties, and diagnosis misclassifications sometimes occur. Hence, some AD individuals included in enriched AD endophenotypes may present concomitant vascular brain disease. The generation of large histopathological GWAS cohorts with associated quantitative data on each pathological hallmark is the ultimate solution to tackling the intrinsic pathologic heterogeneity observed in AD dementia. Unfortunately, there are few examples of neuropathological cohorts: only one GWAS has investigated the genetics of CAA, being *APOE* the unique GWS signal.^24^ Furthermore, in this study, small number of AD cases evolved to vascular dementia during follow-up. Large clinical GWAS cannot control diagnostic changes occurring in clinical practice. Clinical diagnosis is a dynamic variable, so understanding the genetic profiles of specific subgroups of patients that develop other pathologies would provide relevant and powerful information. Finally, the exact effector genes for LOAD genetic findings remain unclear. This is a severe limitation to pathway analysis that can only be circumvented by isolating the causative mutations. Independent replication will be needed to corroborate our new reported GWS signals. In that sense, the selection of specific patient groups might lead to successful replication studies.

The assessment of heterogeneity has important implications for gene discovery, the development of treatments and their appropriate use in individual patients. In that sense, the GR@ACE cohort provides useful genomic information, as it accounts for potential sources of variability and contains different sub-groups of cases. This enabled us to analyze the LOAD genetic landscape in terms of clinical endophenotypes. Our efforts to disentangle the mechanistic pathways operating under clinical sub-groups of patients revealed that vasculature regulation may be an essential part of the causative mechanism of LOAD. Finally, our exploration of AD genetics highlights the relevance of sample composition in genetic discoveries. Considering sample composition in the design of genetic studies might lead to the identification of genetic profiles, which can help clinicians distinguish subsets of patients within the disease spectrum and promote novel therapy targets for Alzheimer’s disease.

## Supporting information

Appendix_Section

## Acknowledgements

We would like to thank patients and controls who participated in this project. The Genome Research@Fundació ACE project (GR@ACE) is supported by Fundación bancaria “La Caixa”, Grifols SA, Fundació ACE and ISCIII. We also want to thank the private sponsors who support the basic and clinical projects of our institution (Piramal AG, Laboratorios Echevarne, Araclon Biotech S.A. and Fundació ACE). We are indebted to the Trinitat Port-Carbó legacy and her family for their support of Fundació ACE research programs. Fundació ACE is a participating center in the Dementia Genetics Spanish Consortium (DEGESCO). A.R. and M.B. receive support from the European Union/EFPIA Innovative Medicines Initiative Joint Undertaking ADAPTED and MOPEAD projects (Grants No. 115975 and 115985, respectively). M.B. and A.R. are also supported by national grants PI13/02434, PI16/01861 and PI17/01474. Acción Estratégica en Salud is integrated into the Spanish National R + D + I Plan and funded by ISCIII (Instituto de Salud Carlos III)-Subdirección General de Evaluación and the Fondo Europeo de Desarrollo Regional (FEDER- “Una manera de Hacer Europa”). LMR is supported by Consejería de Salud de la Junta de Andalucía (Grant PI-0001/2017) Control samples and data from patients included in this study were provided in part by the National DNA Bank Carlos III (www.bancoadn.org, University of Salamanca, Spain) and Hospital Universitario Virgen de Valme (Sevilla, Spain); they were processed following standard operating procedures with the appropriate approval of the Ethical and Scientific Committee. The present work was performed as part of the Biochemistry, Molecular Biology and Biomedicine doctoral program of S. Moreno-Grau at Universitat Autònoma de Barcelona (Barcelona, Spain).

Data collection and sharing for this project was partially funded by the **Alzheimer’s Disease Neuroimaging Initiative (ADNI)** (National Institutes of Health Grant U01 AG024904) and DOD ADNI (Department of Defense award number W81XWH-12-2-0012). ADNI is funded by the National Institute on Aging and the National Institute of Biomedical Imaging and Bioengineering, as well as through generous contributions from the following: AbbVie; the Alzheimer’s Association; the Alzheimer’s Drug Discovery Foundation; Araclon Biotech; BioClinica, Inc.; Biogen; Bristol-Myers Squibb Company; CereSpir, Inc.; Cogstate; Eisai Inc.; Elan Pharmaceuticals, Inc.; Eli Lilly and Company; EuroImmun; F. Hoffmann-La Roche Ltd and its affiliated company Genentech, Inc.; Fujirebio; GE Healthcare; IXICO Ltd.; Janssen Alzheimer Immunotherapy Research & Development, LLC.; Johnson & Johnson Pharmaceutical Research & Development LLC.; Lumosity; Lundbeck; Merck & Co., Inc.; Meso Scale Diagnostics, LLC.; NeuroRx Research; Neurotrack Technologies; Novartis Pharmaceuticals Corporation; Pfizer Inc.; Piramal Imaging; Servier; Takeda Pharmaceutical Company; and Transition Therapeutics. The Canadian Institutes of Health Research provide funds to support ADNI clinical sites in Canada. Private sector contributions are facilitated by the Foundation for the National Institutes of Health (www.fnih.org). The grantee organization is the Northern California Institute for Research and Education, and the study was coordinated by the Alzheimer’s Therapeutic Research Institute at the University of Southern California. ADNI data are disseminated by the Laboratory for Neuro Imaging at the University of Southern California.

The **AddNeuroMed** data are from a public-private partnership supported by EFPIA companies and SMEs as part of InnoMed (Innovative Medicines in Europe), an Integrated Project funded by the European Union of the Sixth Framework program priority FP6-2004-LIFESCIHEALTH-5. Clinical leads responsible for data collection are Iwona Kłoszewska (Lodz), Simon Lovestone (London), Patrizia Mecocci (Perugia), Hilkka Soininen (Kuopio), Magda Tsolaki (Thessaloniki) and Bruno Vellas (Toulouse). Imaging leads are Andy Simmons (London), Lars-Olad Wahlund (Stockholm) and Christian Spenger (Zurich). Bioinformatics leads are Richard Dobson (London) and Stephen Newhouse (London).

Funding support for the **Alzheimer’s Disease Genetics Consortium (ADGC)** was provided through the NIA Division of Neuroscience (U01-AG032984).

The genotypic and associated phenotypic data used in the study “Multi-Site Collaborative Study for Genotype-Phenotype Associations in Alzheimer’s Disease (**GenADA**)” were provided by GlaxoSmithKline, R&D Limited. The datasets used for the analyses described in this manuscript were obtained from dbGaP at http://www.ncbi.nlm.nih.gov/gap through dbGaP accession number phs000219.

The **Mayo Clinic Alzheimer’s Disease Genetic Studies**, led by Dr. Nilüfer Ertekin-Taner and Dr. Steven G. Younkin at the Mayo Clinic in Jacksonville, FL, used samples from the Mayo Clinic Study of Aging, the Mayo Clinic Alzheimer’s Disease Research Center and the Mayo Clinic Brain Bank. Data collection was supported through funding by NIA grants P50 AG016574, R01 AG032990, U01 AG046139, R01 AG018023, U01 AG006576, U01 AG006786, R01 AG025711, R01 AG017216 and R01 AG003949, NINDS grant R01 NS080820, the CurePSP Foundation and support from the Mayo Foundation.

The **Neocodex-Murcia** study was funded by the Fundación Alzheimur (Murcia), the Ministerio de Educación y Ciencia (Gobierno de España), Corporación Tecnológica de Andalucía, Agencia IDEA (Consejería de Innovación, Junta de Andalucía), the Diabetes Research Laboratory and the Biomedical Research Foundation. University Hospital Clínico San Carlos has been supported by CIBER de *Diabetes y Enfermedades Metabólicas Asociadas* (CIBERDEM); CIBERDEM is an ISCIII Project.

The **ROS/MAP** study data were provided by the Rush Alzheimer’s Disease Center, Rush University Medical Center, Chicago. Data collection was supported through funding by NIA grants P30AG10161, R01AG15819, R01AG17917, R01AG30146, R01AG36836, U01AG32984 and U01AG46152, the Illinois Department of Public Health and the Translational Genomics Research Institute

The **TGEN** study was supported by Kronos Life Sciences Laboratories, the National Institute on Aging (Arizona Alzheimer’s Disease Center grants P30 AG19610 and RO1 AG023193, the Mayo Clinic Alzheimer’s Disease Center grant P50 AG16574 and the Intramural Research Program), the National Alzheimer’s Coordinating Center (U01 AG016976) and the state of Arizona.

***The results published here are in part based on data obtained from the AMP-AD Knowledge Portal accessed at http://dx.doi.org/doi:10.7303/syn2580853***

## References

1 Baumgart M, Snyder HM, Carrillo MC, Fazio S, Kim H, Johns H. Summary of the evidence on modifiable risk factors for cognitive decline and dementia: A population-based perspective. Alzheimer’s Dement 2015; 11: 718–26.

2 Brookmeyer R, Johnson E, Ziegler-Graham K, Arrighi HM. Forecasting the global burden of Alzheimer’s disease. Alzheimer’s Dement 2007; 3: 186–91.

3 Serrano-Pozo A, Frosch MP, Masliah E, Hyman BT. Neuropathological Alterations in Alzheimer Disease. Cold Spring Harb Perspect Med 2011; 1: a006189–a006189.

4 Boyle PA, Yu L, Leurgans SE, et al. Attributable Risk of Alzheimer’s Dementia Due to Age-Related Neuropathologies. Ann Neurol 2018; published online Nov 12. DOI:10.1002/ana.25380.

5 Viswanathan A, Rocca WA, Tzourio C. Vascular risk factors and dementia: How to move forward? Neurology 2009; 72: 368–74.

6 Goate A, Chartier-Harlin MC, Mullan M, et al. Segregation of a missense mutation in the amyloid precursor protein gene with familial Alzheimer’s disease. Nature 1991; 349: 704–6.

7 Sherrington R, Rogaev EI, Liang Y, et al. Cloning of a gene bearing missense mutations in early-onset familial Alzheimer’s disease. Nature 1995; 375: 754–60.

8 Sherrington R, Froelich S, Sorbi S, et al. Alzheimer’s disease associated with mutations in presenilin 2 is rare and variably penetrant. Hum Mol Genet 1996; 5: 985–8.

9 Brainstorm Consortium V, Anttila V, Bulik-Sullivan B, et al. Analysis of shared heritability in common disorders of the brain. Science 2018; 360: eaap8757.

10 Avramopoulos D. Genetics of Alzheimer’s disease: recent advances. Genome Med 2009; 1: 34.

11 Corder E, Saunders A. Gene dose of apolipoprotein E type 4 allele and the risk of Alzheimer’s disease in late onset families. Science (80-) 1993; 8: 41–3.

12 Kunkle BW, Grenier-Boley B, Sims R, et al. Meta-analysis of genetic association with diagnosed Alzheimer’s disease identifies novel risk loci and implicates Abeta, Tau, immunity and lipid processing. bioRxiv 2018;: 294629.

13 Sims R, van der Lee SJ, Naj AC, et al. Rare coding variants in PLCG2, ABI3, and TREM2 implicate microglial-mediated innate immunity in Alzheimer’s disease. Nat Genet 2017; 49: 1373–84.

14 Jansen I, Savage J, Watanabe K, et al. Genetic meta-analysis identifies 10 novel loci and functional pathways for Alzheimer’s disease risk. bioRxiv 2018;: 258533.

15 Ridge PG, Hoyt KB, Boehme K, et al. Assessment of the genetic variance of late-onset Alzheimer’s disease. Neurobiol Aging 2016; 41: 200.e13–200.e20.

16 Boada M, Tárraga L, Hernández I, et al. Design of a comprehensive Alzheimer’s disease clinic and research center in Spain to meet critical patient and family needs. Alzheimers Dement 2014; 10: 409–15.

17 Román GC, Tatemichi TK, Erkinjuntti T, et al. Vascular dementia: diagnostic criteria for research studies. Report of the NINDS-AIREN International Workshop. Neurology 1993; 43: 250–60.

18 Lambert JC, Ibrahim-Verbaas CA, Harold D, et al. Meta-analysis of 74,046 individuals identifies 11 new susceptibility loci for Alzheimer’s disease. Nat Genet 2013; 45: 1452–8.

19 Jun GR, Chung J, Mez J, et al. Transethnic genome-wide scan identifies novel Alzheimer’s disease loci. Alzheimers Dement 2017; 13: 727–38.

20 Regan C, Katona C, Walker Z, Hooper J, Donovan J, Livingston G. Relationship of vascular risk to the progression of Alzheimer disease. Neurology 2006; 67: 1357–62.

21 Zhang B, Gaiteri C, Bodea L-G, et al. Integrated systems approach identifies genetic nodes and networks in late-onset Alzheimer’s disease. Cell 2013; 153: 707–20.

22 Weller RO, Boche D, Nicoll JAR. Microvasculature changes and cerebral amyloid angiopathy in Alzheimer’s disease and their potential impact on therapy. Acta Neuropathol 2009; 118: 87–102.

23 Charidimou A, Shoamanesh A, Al-Shahi Salman R, et al. Cerebral amyloid angiopathy, cerebral microbleeds and implications for anticoagulation decisions: The need for a balanced approach. Int J Stroke 2018; 13: 117–20.

24 Beecham GW, Hamilton K, Naj AC, et al. Genome-wide association meta-analysis of neuropathologic features of Alzheimer’s disease and related dementias. PLoS Genet 2014; 10: e1004606.

25 Biffi A, Shulman JM, Jagiella JM, et al. Genetic variation at CR1 increases risk of cerebral amyloid angiopathy. Neurology 2012; 78: 334–41.

26 Mäkelä M, Kaivola K, Valori M, et al. Alzheimer risk loci and associated neuropathology in a population-based study (Vantaa 85+). Neurol Genet 2018; 4: e211.

27 Sakai K, Yamada M. [Cerebral amyloid angiopathy]. Brain Nerve 2014; 66: 827–35.

28 Villeda SA, Plambeck KE, Middeldorp J, et al. Young blood reverses age-related impairments in cognitive function and synaptic plasticity in mice. Nat Med 2014; 20: 659–63.

29 Boada M, Antúnez C, Ramírez-Lorca R, et al. ATP5H/KCTD2 locus is associated with Alzheimer’s disease risk. Mol Psychiatry 2014; 19: 682–7.

30 Traylor M, Adib-Samii P, Harold D, et al. Shared genetic contribution to Ischaemic Stroke and Alzheimer’s Disease. Ann Neurol 2016; 79: 739.

31 Lucariello M, Vidal E, Vidal S, et al. Whole exome sequencing of Rett syndrome-like patients reveals the mutational diversity of the clinical phenotype. Hum Genet 2016; 135: 1343–54.

32 Hanada K, Kumagai K, Yasuda S, et al. Molecular machinery for non-vesicular trafficking of ceramide. Nature 2003; 426: 803–9.

33 Leduc V, De Beaumont L, Théroux L, et al. HMGCR is a genetic modifier for risk, age of onset and MCI conversion to Alzheimer’s disease in a three cohorts study. Mol Psychiatry 2015; 20: 867–73.

34 Broce IJ, Tan CH, Fan CC, et al. Dissecting the genetic relationship between cardiovascular risk factors and Alzheimer’s disease. Acta Neuropathol 2018; published online Nov 9. DOI:10.1007/s00401-018-1928-6.

35 Escott-Price V, Bellenguez C, Wang L-S, et al. Gene-wide analysis detects two new susceptibility genes for Alzheimer’s disease. PLoS One 2014; 9: e94661.

36 Luo L, Bokil NJ, Wall AA, et al. SCIMP is a transmembrane non-TIR TLR adaptor that promotes proinflammatory cytokine production from macrophages. Nat Commun 2017; 8: 14133.

37 Delunardo F, Margutti P, Pontecorvo S, et al. Screening of a microvascular endothelial cDNA library identifies rabaptin 5 as a novel autoantigen in Alzheimer’s disease. J Neuroimmunol 2007; 192: 105–12.

38 Moreno-Grau SM-G, Hernández I, Heilmann-Heimbach S, et al. Genome-wide significant risk factors on chromosome 19 and the APOE locus. Oncotarget 2018; 9: 24590–600.

