## Supplementary_Figures for "Genome-wide association analysis of dementia and its clinical endophenotypes reveal novel loci associated with Alzheimer’s disease and three causality networks of AD: the GR@ACE project"

### Supplementary Figure 1

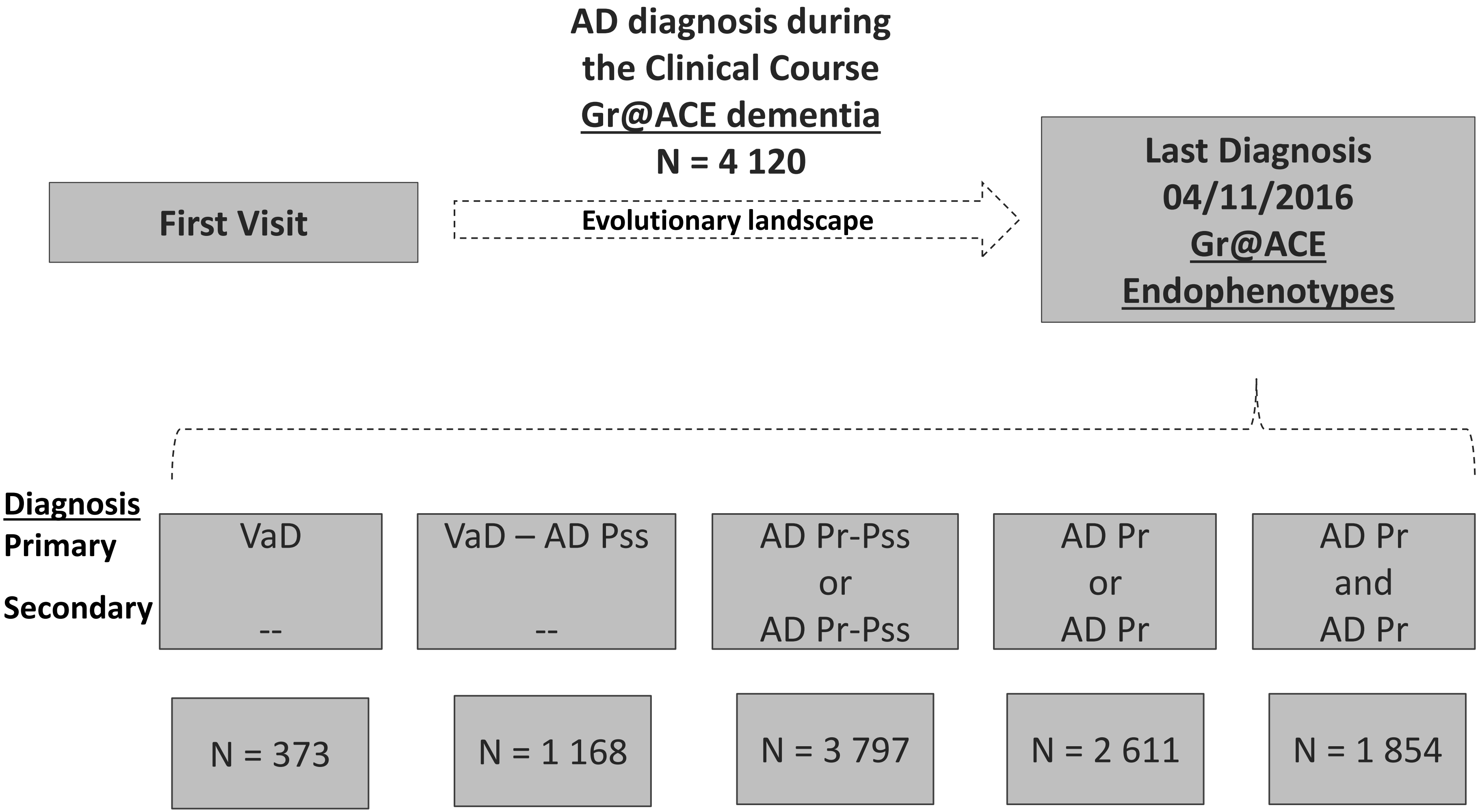

### Supplementary Figure 2

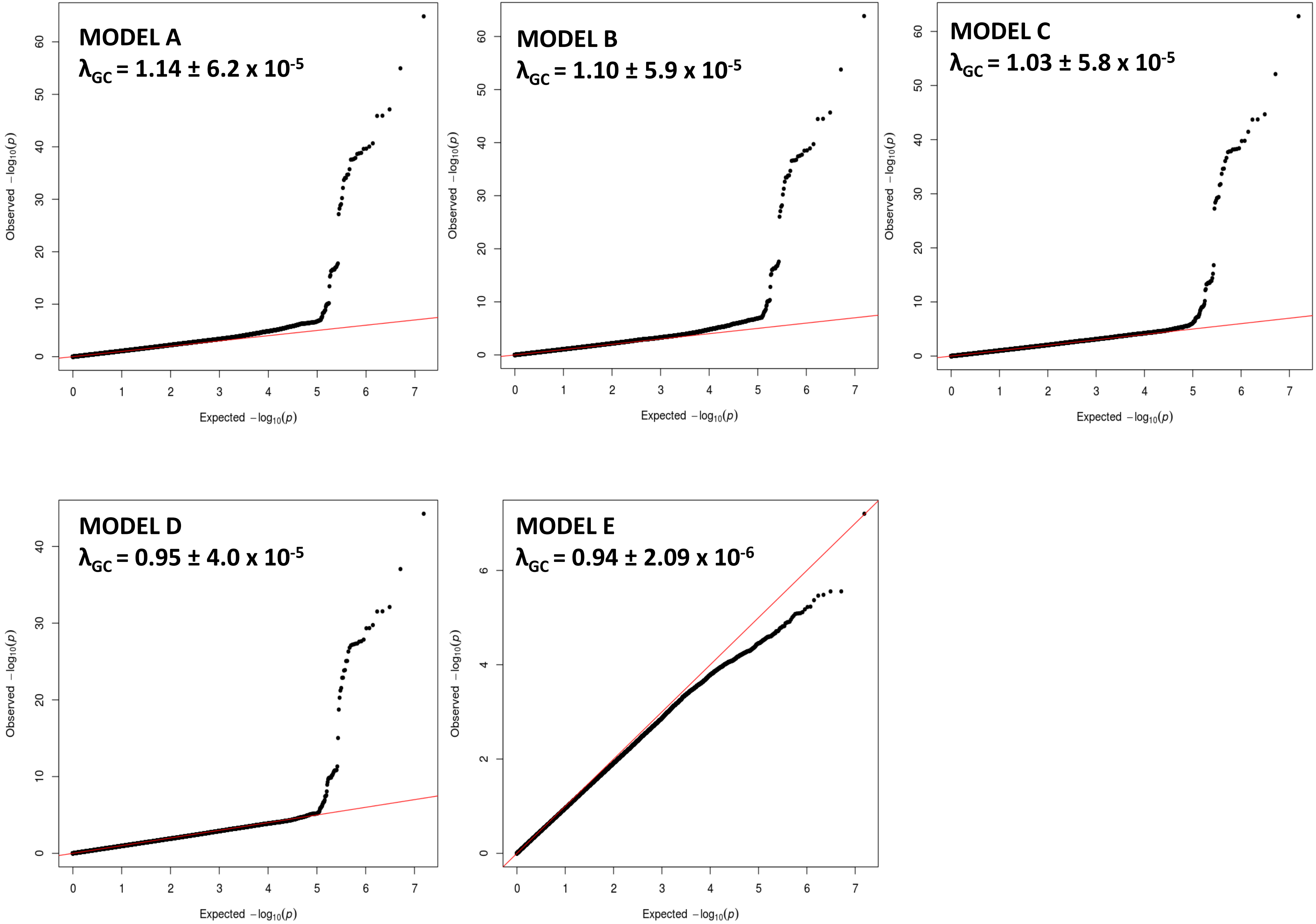

Supplementary Figure 3

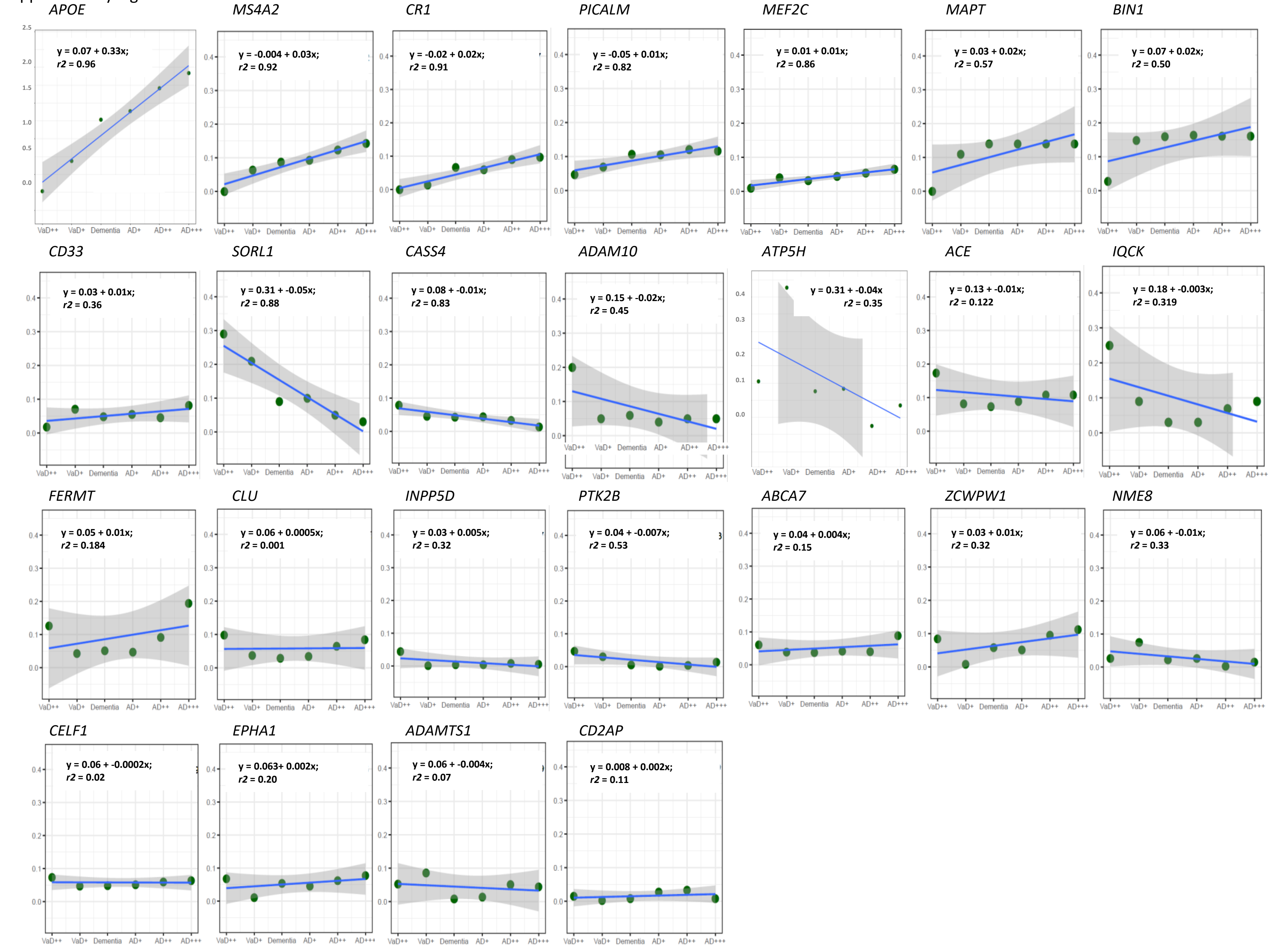

### Supplementary Figure 4

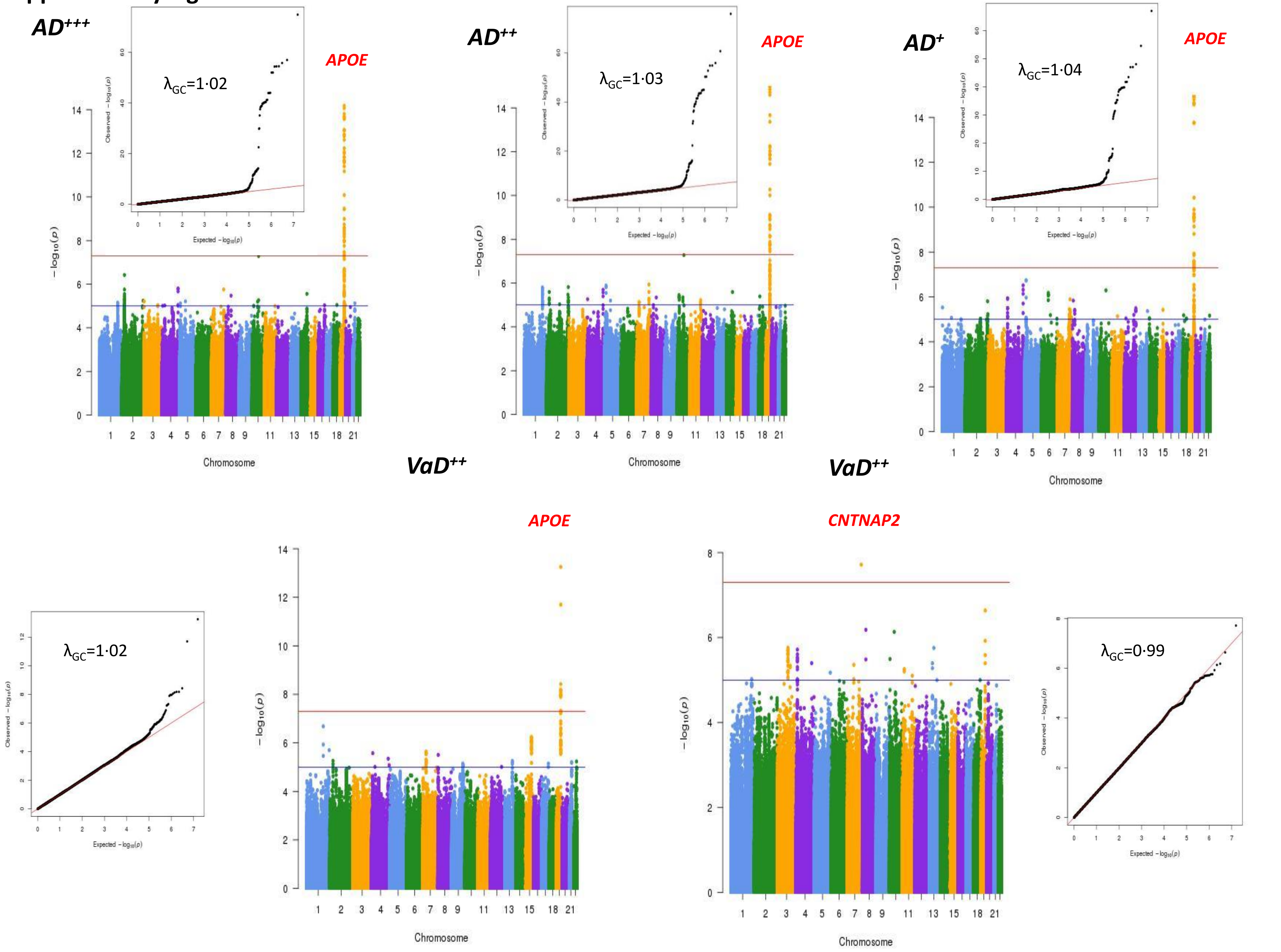

#### Supplementary Figure 5

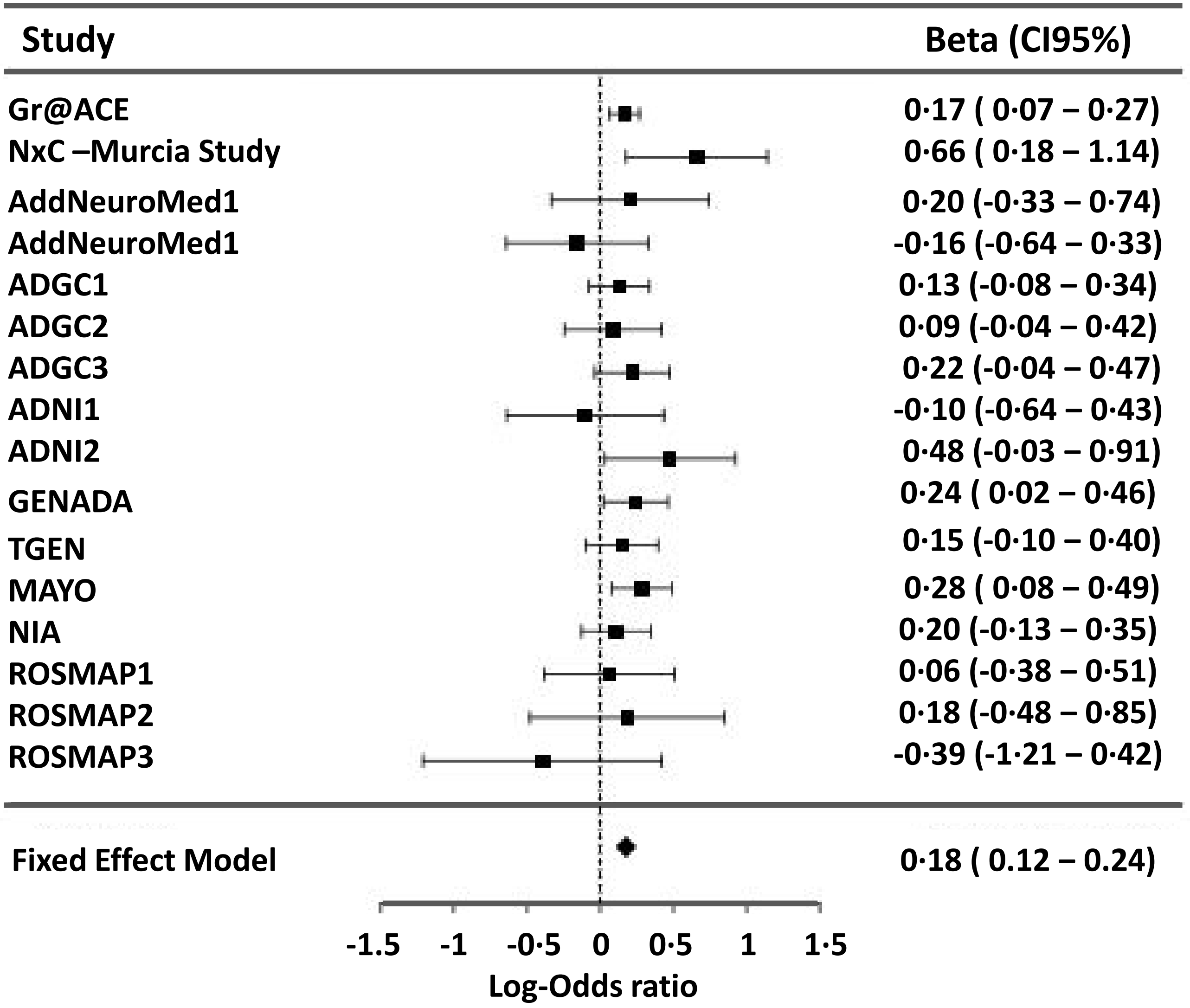

**Supplementary Figure 6**

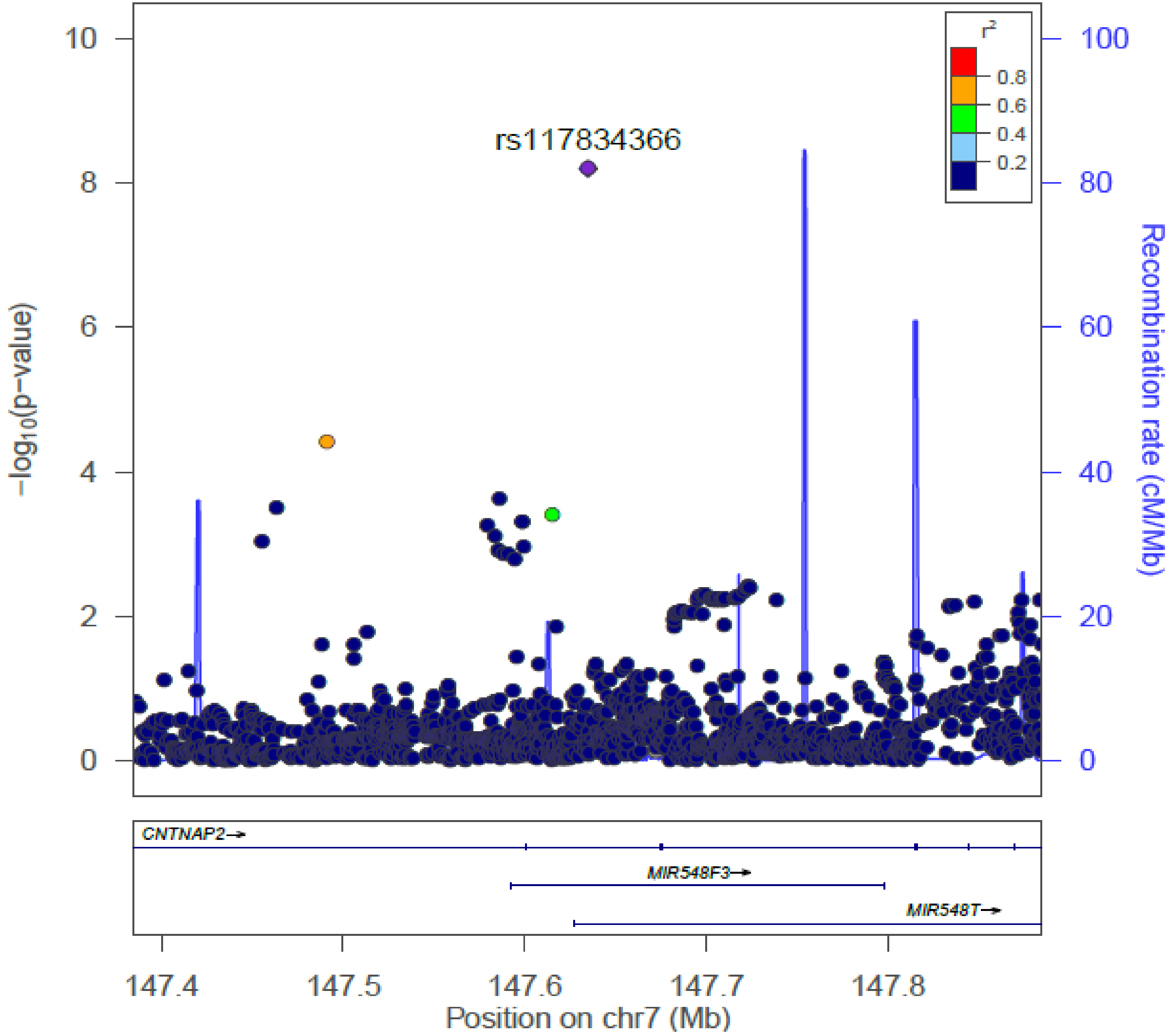

#### Supplementary Figure 7

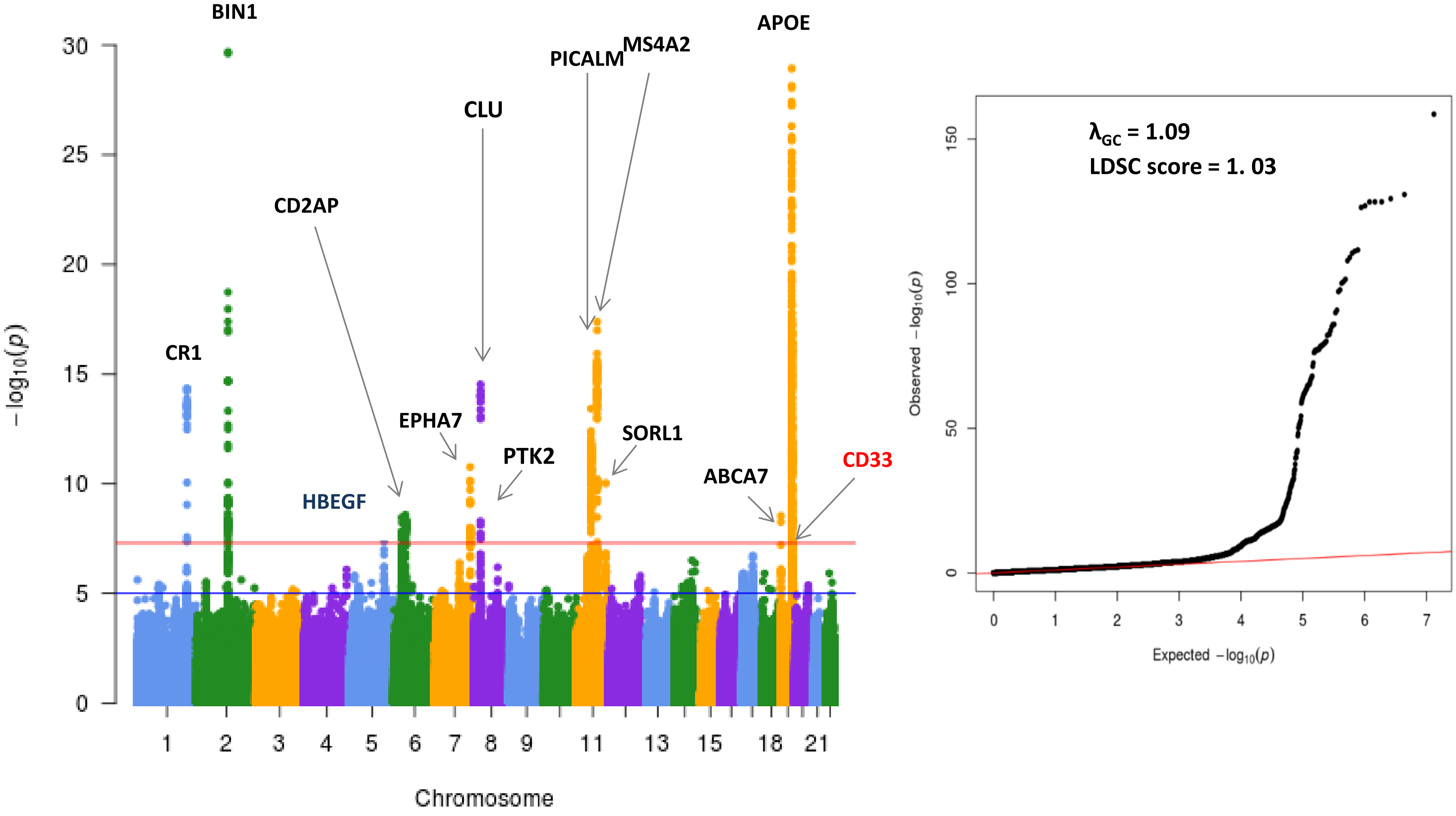

**Supplementary Figure 8**

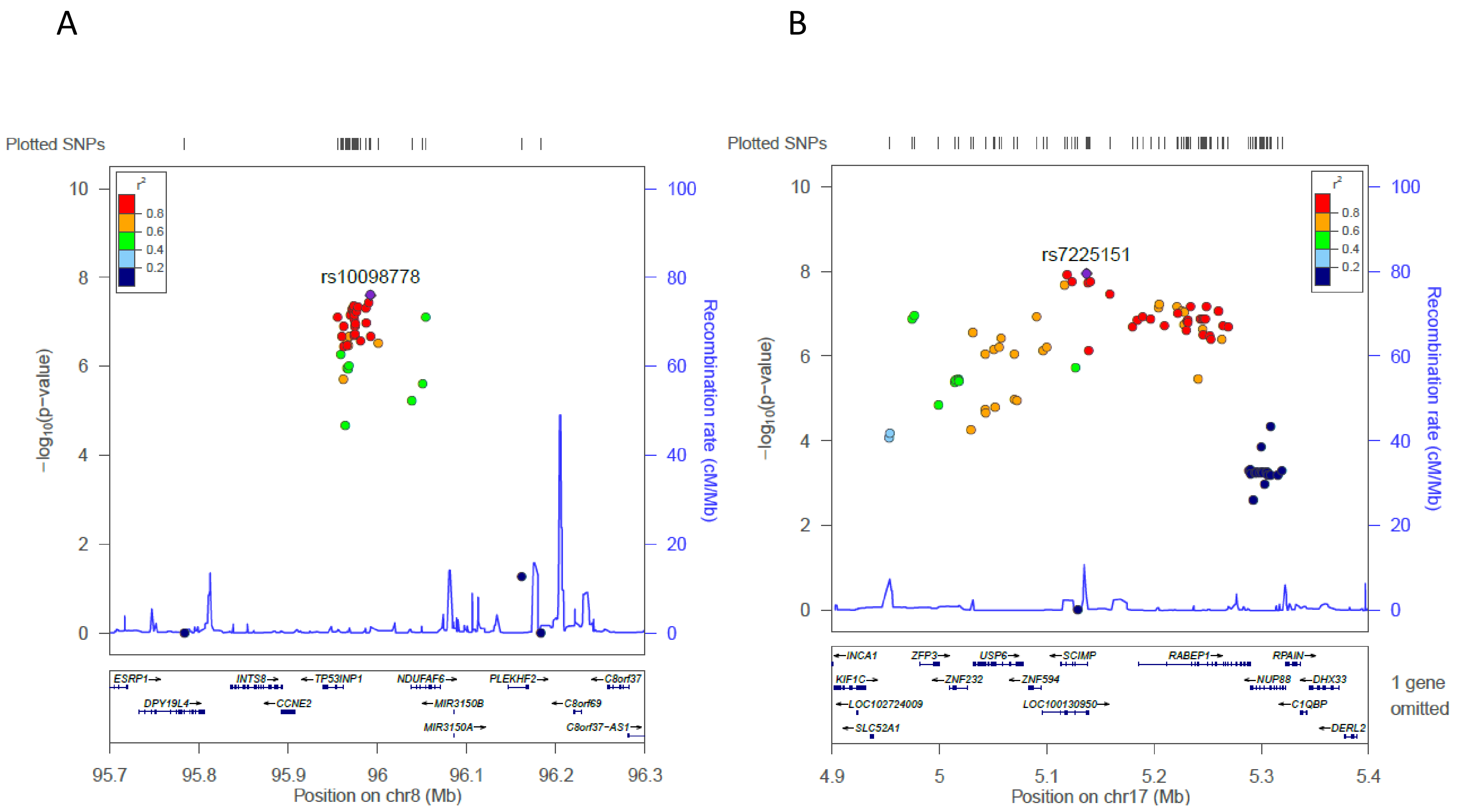
