## Appendix_Section for "Genome-wide association analysis of dementia and its clinical endophenotypes reveal novel loci associated with Alzheimer’s disease and three causality networks of AD: the GR@ACE project"

**Affiliation:**

*21. BT-CIEN*

*22. Department of Molecular Neuropathology, Centro de Biología Molecular "Severo Ochoa" (CBMSO), Consejo Superior de Investigaciones Científicas (CSIC)/Universidad Autónoma de Madrid (UAM)*

*23. Unidad de Demencias. Hospital Clínico Universitario Virgen de la Arrixaca*

*24. Fundación Pública Galega de Medicina Xenómica- CIBERER-IDIS, Santiago de Compostela, Spain.*

****Corresponding author**: Agustín Ruiz M.D. Ph.D.
**Address:** Research Center. Fundació ACE. Institut Català de Neurociències Aplicades. C/ Marquès de Sentmenat, 57. 08029 Barcelona, Spain

**Conflict of Interest:** None.

***The GR@ACE study group***

Abdelnour C[1,2], Aguilera N[1], Alarcon E[1,3], Alegret M[1,2], Boada M[1,2], Buendia M[1], Cañabate P[1,2], Carracedo A[4,5], Corbatón A[6], de Rojas I[1], Diego S[1], Espinosa A[1,2], Gailhajenet A[1], García González P[1], Gil S[1], Guitart M[1], González Pérez A[7], Hernández I[1,2], Ibarria, M[1], Lafuente A[1], Macias J[8], Maroñas O[4], Martín E[1], Martínez MT[6], Mauleón A[1], Monté G[1], Montrreal L[1], Moreno-Grau S[1,2], Orellana A[1], Ortega G[1,2], Pancho A[1], Pelejà E[1], Pérez-Cordon A[1], Pineda JA[8], Preckler S[1], Quintela I[3], Real LM[3,8], Rodríguez-Gómez O[1,2], Rosende-Roca M[1], Ruiz A[1,2], Ruiz S[1,2], Sáez ME[7], Sanabria A[1,2], Santos-Santos MA[1], Serrano-Rios M[6], Sotolongo-Grau O[1], Tárraga L[1,2], Valero S[1,2], Vargas L[1]

*(1. Research Center and Memory clinic Fundació ACE. Institut Català de Neurociències Aplicades. Universitat Internacional de Catalunya. Barcelona. Spain; 2. CIBERNED, Center for Networked Biomedical Research on Neurodegenerative Diseases, National Institute of Health Carlos III, Ministry of Economy and Competitiveness, Spain; 3. Dep. of Surgery, Biochemistry and Molecular Biology, School of Medicine. University of Málaga. Málaga. Spain 4. Grupo de Medicina Xenómica, Centro Nacional de Genotipado (CEGEN-PRB3-ISCIII). Universidade de Santiago de Compostela, Santiago de Compostela, Spain; 5. Fundación Pública Galega de Medicina Xenómica- CIBERER-IDIS, Santiago de Compostela, Spain; 6. Centro de Investigación Biomédica en Red de Diabetes y Enfermedades Metabólicas Asociadas, CIBERDEM, Spain, Hospital Clínico San Carlos, Madrid, Spain 7. CAEBI. Centro Andaluz de Estudios Bioinformáticos., Sevilla, Spain; 8. Unidad Clínica de Enfermedades Infecciosas y Microbiología. Hospital Universitario de Valme, Sevilla, Spain)*

**DEGESCO consortium**

Adarmes-Gómez AD[1,2], Alarcón-Martín E[3,4], Álvarez I[5], Álvarez V[6,7], Amer-Ferrer G[8], Antequera M[9], Antúnez C[9], Baquero M[10], Bernal M[11], Blesa R[2,12], Boada M[2,3], Buiza-Rueda D[1,2], Bullido MJ[2,13,14], Burguera JA[10], Calero M[2,15,16], Carrillo F[1,2], Carrión-Claro M[1,2], Casajeros MJ[17], Clarimón J[2,12], Cruz-Gamero JM[4], de Pancorbo MM[18], de Rojas I[3], del Ser T[14], Diez-Fairen M[5], Fortea J[2,12], Franco E[11], Frank-García A[2,14,19], García-Alberca JM[20], Garcia Madrona S[16], Garcia-Ribas G[16], Gómez-Garre P[1,2], Hernández I[2,3], Hevilla S[20], Jesús S[1,2], Labrador Espinosa MA[1,2], Lage C2,21, Legaz A[9], Lleó A[2,12], López de Munáin A[22], López-García S[2,21], Macias D[1,2], Manzanares S[9,23], Marín M[11], Marín-Muñoz J[9], Marín T[20], Marquié M[3], Martín Montes A[2,13,19], Martínez B[9], Martínez C[7,24], Martínez V[9], Martínez-Lage Álvarez P[25], Medina M[2,14], Mendioroz Iriarte M[26], Menéndez-González M[7,27], Mir P[1,2], Molinuevo JL[28], Monté G[3], Montrreal L[3], Moreno-Grau S[2,3], Orellana A[3], Pastor AB[15], Pastor P[5], Pérez Tur J[2,29,30], Periñán-Tocino T[1,2], Piñol Ripoll G[2,31], Rábano A[2,15,32], Rodrigo S[11], Rodríguez-Rodríguez E[2,21], Royo JL[4], Ruiz A[2,3], Sanchez del Valle Díaz R[33], Sánchez-Juan P[2,21], Sastre I[2,13], Sotolongo-Grau O3, Tárraga L[2,3], Valero S[2,3], Vicente MP[9], Vivancos L[9]

*(1. Unidad de Trastornos del Movimiento, Servicio de Neurología y Neurofisiología. Instituto de Biomedicina de Sevilla (IBiS), Hospital Universitario Virgen del Rocío/CSIC/Universidad de Sevilla, Seville, Spain; 2. CIBERNED, Network Center for Biomedical Research in Neurodegenerative Diseases, National Institute of Health Carlos III, Spain; 3. Research Center and Memory clinic Fundació ACE. Institut Català de Neurociències Aplicades. Universitat Internacional de Catalunya. Barcelona; 4. Dep. of Surgery, Biochemistry and Molecular Biology, School of Medicine. University of Málaga. Málaga. Spain; 5. Fundació per la Recerca Biomèdica i Social Mútua Terrassa, and Memory Disorders Unit, Department of Neurology, Hospital Universitari Mutua de Terrassa, University of Barcelona School of Medicine, Terrassa, Barcelona, Spain; 6. Laboratorio de Genética Hospital Universitario Central de Asturias, Oviedo; 7. Instituto de Investigación Biosanitaria del Principado de Asturias (ISPA); 8. Department of Neurology, Hospital Universitario Son Espases, Palma, Spain; 9. Unidad de Demencias. Hospital Clínico Universitario Virgen de la Arrixaca; 10. Servei de Neurologia, Hospital Universitari i Politècnic La Fe; 11. Unidad de Demencias, Servicio de Neurología y Neurofisiología. Instituto de Biomedicina de Sevilla (IBiS), Hospital Universitario Virgen del Rocío/CSIC/Universidad de Sevilla, Seville, Spain; 12. Memory Unit, Neurology Department and Sant Pau Biomedical Research Institute, Hospital de la Santa Creu i Sant Pau, Universitat Autònoma de Barcelona, Barcelona, Spain; 13. Centro de Biologia Molecular Severo Ochoa (C.S.I.C.-U.A.M.), Universidad Autonoma de Madrid, Madrid, Spain; 14. Instituto de Investigacion Sanitaria 'Hospital la Paz' (IdIPaz), Madrid, Spain; 15. CIEN Foundation, Queen Sofia Foundation Alzheimer Center, Madrid, Spain; 16. Instituto de Salud Carlos III (ISCIII), Madrid, Spain; 17. Hospital Universitario Ramón y Cajal; Madrid, Spain; 18. BIOMICs, País Vasco; Centro de Investigación Lascaray. Universidad del País Vasco UPV/EHU; 19. Neurology Service, Hospital Universitario La Paz (UAM), Madrid, Spain; 20. Alzheimer Research Center & Memory Clinic. Andalusian Institute for Neuroscience. Málaga. Spain; 21. Neurology Service, Marqués de Valdecilla University Hospital (University of Cantabria and IDIVAL), Santander, Spain; 22. Hospital Donostia de San Sebastían; 23. Fundación para la Formación e Investigación Sanitarias de la Región de Murcia; 24. Servicio de Neurología -Hospital de Cabueñes-Gijón; 25. Centro de Investigación y Terapias Avanzadas. Fundación CITA-alzheimer; 26. Navarrabiomed; 27. Servicio de Neurología -Hospital Universitario Central de Asturias, Oviedo; 28. Barcelona βeta Brain Research Center – Fundació Pasqual Maragall; 29. Unitat de Genètica Molecular. Institut de Biomedicina de València-CSIC; 30. Unidad Mixta de Neurologia Genètica. Instituto de Investigación Sanitaria La Fe; 31. Unitat Trastorns Cognitius, Hospital Universitari Santa Maria de Lleida, Institut de Recerca Biomédica de Lleida (IRBLLeida), Lleida, España. 32. BT-CIEN; 33. Hospital Clínic Barcelona.)*

**Alzheimer's Disease Neuroimaging Initiative**: Data used in preparing this article were obtained from the Alzheimer's Disease Neuroimaging Initiative (ADNI) database (adni.loni.usc.edu). As such, the investigators within the ADNI contributed to the design and implementation of ADNI and/or provided data but did not participate in the analysis or writing of this report. A complete listing of ADNI investigators can be found at http://adni.loni.usc.edu/wp-content/uploads/how_to_apply/ADNI_Acknowledgement_List.pdf

**Supplementary Figure Legend.**

Supplementary Figure 1. Flow chart diagram for inclusion of AD patients and construction of GR@ACE clinical endophenotypes.

Supplementary Figure 2. Quantile-quantile plots across five regression models using the GR@ACE cohort.

Supplementary Figure 3. Effect change for each individual LOAD genetic variant across GR@ACE clinical endophenotypes.

Supplementary Figure 4. A) Results of genome-wide association analysis for GR@ACE clinical endophenotypes. B) Quantile-quantile plot.

Supplementary Figure 5. Associations of the region centered on rs117834366 located in the *CNTNAP2* gene.

Supplementary Figure 6. Forest plot for the effect of rs4704171-*ANKDR31* in risk of AD.

Supplementary Figure 7. Results of genome-wide association analysis for GR@ACE meta-analysis with IGAP Stage I. B) QQplot.

Supplementary Figure 8. Associations of the region centered on rs10098778 located in the *TP53INP1/NDUFAF6* gene. B) on rs7225151 located in the *SCIMP* gene.

**Supplementary Tables Legend.**

Supplementary Table 1. Association results obtained from the GR@ACE dataset and GR@ACE endophenotypes, classified per gene categories for functional pathway analysis. Global effect change per genetic variant.

Supplementary Table 2. Full description of dbGaP datasets used in the meta-analysis.

Supplementary Table 3. eQTL analysis for novel GWAS significant hits associated with AD.

Supplementary Table 4. Association results for rs2732703-*KANSL1*/*MAPT* in *APOE* ε4 carriers and non-carriers across GR@ACE endophenotypes.

Supplementary Table 5. Replication results for rs7100488-*PCBD1/UNSC5B* and rs117834366-*CNTNAP2* in additional datasets.

Supplementary Table 6. Top ten biological pathways per gene cluster after sub-analysis of gene Category C.

Supplementary Table 7. Top ten biological pathways per gene cluster after secondary gene categorization.

Supplementary Table 8. Association results for known LOAD loci in meta-analysis with dbGaP datasets and IGAP Stage I and Stage I+II.

### Methods

#### GR@ACE cohort and phenotype definitions

All AD patients included in the GR@ACE study received a thorough structured neurological evaluation that included: history, examination, Mini-Mental State Examination (MMSE)[1], Blessed Dementia Rating Scale (BDRS)[2,3], Neuropsychiatric Inventory-questionnaire (NPI-Q)[4], Tinnetti scale for gait and balance[5], Clinical Dementia Rating (CDR)[6], Global Deterioration Scale (GDR)[7] scoring and Hachinski Ischemia Scale[8]. The Fundacio ACE neuropsychological battery (NBACE)[9] was administered to all patients. The NBACE includes measures of cognitive information processing speed, orientation, attention, verbal learning and memory, language, visuoperception, praxis and executive functions. Family members or caregivers are interviewed by a social worker.

Endophenotype approach was feasible due to Fundació ACE's endorsement both, a primary and a secondary diagnosis, as well as routine follow-up evaluations. The secondary diagnosis might be the same or an additional clinical condition that explains the clinical symptoms. Follow-up visits enabled controlling for the progression of the disease and identifying comorbidities and sometimes diagnosis changes, thereby providing a longitudinal landscape for each individual's clinical evolution (Supplementary figure 1). VaD patients were defined according to NINDS-AIREN criteria.[10]

#### GWAS genotyping

Peripheral blood was taken from all individuals to isolate germline DNA from leukocytes. DNA extraction was performed automatically according to standard procedures using the Chemagic system (Perkin Elmer). Extensive DNA quality control was conducted. Only samples reaching DNA concentrations >10ng/µl and presenting high integrity were included for genotyping. Samples and controls were randomized across sample plates to avoid batch effects.

The genotyping array, Axiom 815K Spanish Biobank array, is an adaptation of the Axiom Biobank Genotyping Array, but also contains population-specific rare variations observed in the Spanish population. DNA samples were genotyped according to the manufacturer's instructions (Axiom™ 2.0 Assay Manual Workflow). The Axiom 2.0 assay interrogates biallelic SNPs and simple indels in a single assay workflow. Starting with 200 ng of genomic DNA, the samples were processed through a manual target preparation protocol followed by automated processing of the array plates in the GeneTitan Multi-Channel (MC) instrument. Target preparation involved DNA amplification, fragmentation, purification and resuspension of the target in a hybridization cocktail. The hyb-ready targets were then transferred to the GeneTitan MC instrument for automated, hands-free processing including hybridization, staining, washing and imaging. CEL files were generated using the GeneTitan MC instrument.

To achieve higher genotyping performance, quality control for samples and plates was performed using the Affymetrix power tool (APT) 1.15.0 software following Axiom Data Analysis Workflow. Briefly, sample quality was determined based on the resolution of AT and GC channels in a group of non-polymorphic SNPs (resolution > 0.82). Passing samples were genotyped for sample QC. Samples with a call rate greater than 97% and plates with an average call rate above 98.5% were included for final SNP calling. Quality samples were jointly called to minimize batch effects. Best quality markers were selected for downstream analysis ($N_{SNPs}$ =777,649; 95.4%) using the SNPolisher R package (Thermo Fisher). To assess the sample genotyping concordance, we intentionally re-sampled 200 samples and determined a concordance rate of 99.5%.

### GWAS quality control

Principal component analysis was conducted excluding markers with moderate-to-high linkage disequilibrium (LD) (r2>0.3) and long-range LD regions from PCA analysis using the "indep-pairwise" option of PLINK 1.9[11] (https://www.cog-genomics.org/plink2).

### GWAS Statistical analysis

We tested the association between markers and the targeted phenotypes with and without covariates. We selected a model including the top four PCs as covariates for the discovery stage because this model exhibited the lowest inflation and optimal power compared to alternative models. To establish the discovery model, we conducted an exploratory step using the GR@ACE dataset. We evaluated the performance of five different models: a) Model A, unadjusted; b) Model B, unadjusted and excluding all individuals matching with Hispanic American ancestry; c) Model C, adjusted per four main PCs; d) Model D, adjusted per top four PCs, gender and age; and e) Model E, adjusted per top four PCs, gender, age and *APOE* status (Supplementary figure 2). We explored the QQ plot and the genomic inflation factor ($\lambda$), using the Qqman and GenABEL packages (http://www.genabel.org/packages/GenABEL) from R, to test for covariates in the regression model.

### Genetic exploration of GR@ACE clinical endophenotypes and enrichment analysis

We explored the biological pathways underlying each gene category. First, we extracted the significant co-expressed genes ($p < 0.05$) for each category using GeneFriends in human microarray datasets containing 26,113 experimental conditions and 19,080 genes [12]. Second, to avoid unspecific pathway detection, we selected co-expressed genes with ≥2 LOAD loci per category, and ranked the results from the highest to the lowest number of co-occurrences. Third, we set the maximum number of co-expressed genes for pathway analysis to 150. Thus, *category A* and *category C* includes 150 co-expressed genes (Bonferroni p-value = $3{\cdot}33 \times 10^{-4}$); and *category B*, 99 (Bonferroni p-value = $5{\cdot}05 \times 10^{-4}$). Next, we explored the biological pathways underlying each co-expression on the ranked list by applying an overrepresentation enrichment method in

WebGestalt [13], while using Gene Ontology as a reference database for functional annotations of non-redundant biological pathways.

Next, we performed a sub-classification and pathway analysis for variants from *category C*: Subset C1 includes variants presenting a stable effect across endo-phenotypes (*ABCA7, ADAMTS1, CD2AP, CELF1, EPHA1, INPP5D, NME8, PTK2B, ZCWPW1*); and subset C2 includes variants with an inverse effect between extreme endo-phenotypes (*CLU, FERMT2, IQCK*). Both clusters presented 150 co-expressed genes for pathway analysis.

Finally, we carried out a stringent analysis to validate previous results. We applied a linear regression model using R to evaluate the effect change trend per each genetic variant across clinical endophenotypes. Dementia endophenotypes were the independent variables, coded as follows: $VaD^{++}$ = 1; $VaD^{+}$ = 2; Dementia = 3; $AD^{+++}$ = 6; $AD^{+}$ = 4; $AD^{++}$ = 5. Effect change was the dependent variable. Variants presenting a linear trend a positive effect (*R*2>0.80 and β≥0.01) comprised Category A (*APOE, CR1, MEF2C, MS4A2, PICALM*). Those presenting a linear trend and a negative effect (*R*2>0.80 and β≤-0.01) comprised Category *B* (*SORL1, CASS4*). Those not fulfilling the above criteria comprised Category C (*EPHA1, ABCA7, ACE, ADAMTS1, ADAM10, ATP5H, BIN1, MAPT, CELF1, CD2AP, CD33, CLU, FERMT2, INPP5D, IQCK, NME8, PTK2B, ZCWPW1*). Pathway analysis was conducted applying identical procedure to those described previously. Category A includes 150 co-expressed genes (Bonferroni p-value = $3{\cdot}33 \times 10^{-4}$); Category B, 116 (Bonferroni p-value = $4{\cdot}31 \times 10^{-4}$) and Category C, 150. At this point, we were not able to detect vascular processes in Category A. To discard that this was caused by unspecific pathway detection, we restricted the analysis to include those loci co-expressing with more than 4 LOAD genes at this cluster (co-expressing loci = 43, Bonferroni p-value = $5.55 \times 10^{-4}$). Regulation of vascular development was identified as top pathway ($p < 2{\cdot}14 \times 10^{-7}$).

**Meta-analysis: datasets**

To identify novel loci associated with AD, we combined the GR@ACE dataset and its endophenotypes with: 1) raw genotype data from nine additional GWAS series (N = 21 235) (Supplementary Table 2); and 2) public summary statistics from IGAP stages I (N = 61 571) and II (N = 81 455).

***Genotype level data***

In the first meta-analysis, we had access to raw genotyped data for 7,879 AD patients and 5,947 controls. All datasets were processed by applying the same quality control and imputation procedures as those described for the GR@ACE cohort. To exclude duplicate samples coming from different studies, we also performed a joint analysis and carried out an identity-by-descendent analysis (IBD) (Pi-hat > 0.80) of the nine cohorts and GR@CE (n = 21,235) using PLINK software 1.9.[11] The study cohorts included:

*The Alzheimer's Disease Neuroimaging Initiative (ADNI).*

We obtained the data used in preparing this article from the Alzheimer's Disease Neuroimaging Initiative (ADNI) database (adni.loni.usc.edu). The ADNI was launched in 2003 as a public-private partnership led by Principal Investigator Michael W. Weiner, MD. The primary goal of ADNI is to test whether serial magnetic resonance imaging (MRI), positron emission tomography (PET), other biological markers and clinical and neuropsychological assessments can be combined to measure the progression of mild cognitive impairment (MCI) and early Alzheimer's disease (AD). The ADNI study has three phases: ADNI1, ADNI GO and ADNI2. For up-to-date information, see www.adni-info.org. In the present study, we included 478 cases and 243 controls from ADNI1 and ADNI2.

*The AddNeuroMed Study.*

AddNeuroMed was a public-private partnership for biomarker discovery and replication in Alzheimer's disease [14,15]. It was a multi-center study in Europe with the first patient enrolled in January 2006 and the last in February 2008. The study protocol was planned for a baseline assessment visit with follow-ups every 3 months for the first year, then annual visits that continued through 2013. The study enrolled a total of 258 AD, 257 MCI and 266 controls, but not all had complete data at each assessment. In the present study, we included 450 cases and 187 controls.

This dataset was downloaded from Synapse (doi:10.7303/syn2790911).

*The Alzheimer's Disease Genetics Consortium (ADGC).*

The National Institute on Aging (NIA) Alzheimer's Disease Centers' (ADC) cohort includes subjects ascertained and evaluated by the clinical and neuropathology cores of the 29 NIA-funded ADCs [16]. Data collection was coordinated by the National Alzheimer's Coordinating Center (NACC). The ADC cohort consists of autopsy-confirmed and clinically-confirmed AD cases, cognitively normal elders (CNEs) with complete neuropathology data who were older than 60 years at age of death, as well as living CNEs evaluated using the Uniform dataset (UDS) protocol who were documented to not have mild cognitive impairment (MCI) and were between 60 and 100 years of age at assessment. In the present study, we included 3287 cases and 1322 controls.

This study was downloaded from dbGaP (phs000372).

*Multi-Site Collaborative Study for Genotype-Phenotype Associations in Alzheimer's disease and Longitudinal follow-up of Genotype-Phenotype Associations in Alzheimer's disease and Neuroimaging component of Genotype-Phenotype Associations in Alzheimer's disease (GenADA).*

GenADA was a multi-site collaborative study involving GlaxoSmithKline Inc and nine medical centers in Canada, including 1000 AD patients and 1000 ethnically-matched controls in order to associate DNA sequence (allelic) variations in candidate genes with AD phenotypes [17,18]. The study consisted of both retrospective and prospective data. Where possible, biological relatives with Alzheimer's (up to third-degree relationships such as cousins) and unaffected siblings of AD cases were also recruited. In the present study, we included 785 cases and 764 controls.

This study was downloaded from dbGaP (phs000219).

*The Mayo Clinic LOAD genome-wide association study.*

Subjects from the Mayo LOAD GWAS were selected from two clinical AD Case-Control series: Mayo Clinic Jacksonville (MCJ) and Mayo Clinic Rochester (MCR), as well as a neuropathological series of autopsy-confirmed subjects from the Mayo Clinic Brain Bank [19]. All subjects from the clinical series (MCJ and MCR) were diagnosed by a Mayo Clinic neurologist; all control subjects had a Clinical Dementia Rating score of zero at the most recent time of testing; all LOAD patients had a diagnosis of probable or possible AD according to the NINCDS-ADRDA criteria [20]. All ADs had definite diagnoses according to the NINCDS-ADRDA criteria and had Braak scores of ≥4.0. All non-AD Controls had Braak scores of ≤2.5; many had brain pathology unrelated to AD. . In the present study, we included 703 cases and 1066 controls.

This dataset was downloaded from Synapse (doi:10.7303/syn5550404).

*The Neocodex-Murcia study.*

The study included 327 sporadic AD patients and 801 controls with unknown cognitive status from the Spanish general population collected by Neocodex [21,22]. AD patients were diagnosed as possible or probable AD in accordance with the criteria of the National Institute of Neurological and Communicative Disorders and Stroke and the Alzheimer's Disease and Related Disorders Association (NINCDS-ADRDA)[20]. In the present study, we included 324 cases and 754 controls.

*The Religious Orders Study and Memory and Aging Project (ROS/MAP) Study.*

The Religious Orders Study (ROS) was a longitudinal clinical-pathologic cohort study of aging and Alzheimer's disease (AD) from Rush University that enrolled individuals from religious communities for longitudinal clinical analysis and brain donation [23]. Participants were enrolled from more than 40 groups of religious orders (nuns, priests, brothers) across the United States. Enrolment required no known indications of dementia. Medical conditions were documented starting in 1994 by clinical evaluation or self-report. Alzheimer's disease status was determined by a computer algorithm based on cognitive test performance with a series of discrete clinical judgments made in series by a neuropsychologist and a clinician.

The Memory and Aging Project (MAP) was a longitudinal epidemiologic clinical-pathologic cohort study of common chronic conditions of aging with an emphasis on declines in cognitive and motor function and the risk of Alzheimer’s disease. This study began in 1997 and was run by Rush University [23]. This study was designed to complement the ROS study by enrolling individuals with a wider range of life experiences and socioeconomic status into a study of similar structure and design as ROS. The study enrolled older individuals without any signs of dementia, primarily recruiting from continuous care retirement communities throughout northeastern Illinois, USA. Diagnoses of dementia and AD were performed in an identical manner to the ROS study. . In the present study, we included 628 cases and 229 controls.

This dataset was downloaded from Synapse (doi:10.7303/syn3219045).

*The Translational Genomics Research Institute (TGEN) study.*

The TGEN GWAS study included 643 late-onset AD cases and 404 controls from a neuropathological cohort, and 197 late-onset AD cases and 114 controls from a clinical cohort, all of which were genotyped with the Affimetrix 500 K GeneChip Array [24]. In the present study, we included 741 cases and 449 controls.

TGEN investigators have provided free access to genotype data to other researchers via Coriell Biorepositories (http://www.coriell.org/).

***IGAP summary statistics***

In the second meta-analysis, we combined public summary statistics from IGAP stages I and II (http://web.pasteur-lille.fr/en/recherche/u744/igap/igap_download.php) with the GR@ACE dataset and its endophenotypes. IGAP stage I consisted of 17,008 AD cases and 37,154 controls collected from four published GWAS datasets (The European Alzheimer's disease Initiative – EADI; the Alzheimer Disease Genetics Consortium – ADGC; the Cohorts for Heart and Aging Research in Genomic Epidemiology consortium – CHARGE; the Genetic and Environmental Risk in AD consortium – GERAD) containing 7,055,881 single nucleotide polymorphisms (SNPs). IGAP stage II was a replication effort in which significantly associated variants of stage I were genotyped ($N_{SNP}$ = 11,632) in 8,572 AD cases and 11,312 controls.

**Meta-analysis: association analysis and biological interpretation.**

Summary statistics for each individual dataset were combined with METAL software [25], release 2011-03-25, using inverse variance weighted meta-analysis without genomic control as default. Meta-analysis with genomic control was also explored. To define independent significant signals in the meta-analyses, we assigned variants to clusters using the clump function of PLINK software 1.9[11] based on the association p-values and the short-range LD (250Kb). We used GCTA-COJO[26], version 1.91.3 beta3, to perform standard conditional analysis adjusting for the lead SNP in a region of ±500kb. We estimated the LD score intercept using LD score regression (LDSC v.1.0.0) to distinguish polygenicity from other confounding factors.[27] Regional plots were generated using LocusZoom software.[28]

Finally, we conducted gene expression quantitative trait locus (eQTL) analysis to link GWAS top signals to genes. We used brain (n = 11) and whole blood (n = 1) tissues from the GTEx repository (https://gtexportal.org/home/) for mapping cis-eQTLs. Markers in moderate-to-high LD ($r^2 \geq 0.6$) with the novel lead markers were identified using LDlink[29] and were included in this analysis (Supplementary Table 3).

### Results

#### GR@ACE genome-wide association study

Genome-wide analysis for GR@ACE clinical endophenotypes revealed one novel variant reaching GWS in the VaD$^{++}$ endophenotypes [*CNTNAP2*-rs117834366; OR = 6·03 (3·22 – 11·30); p = 1·91 x $10^{-8}$] (table 3) and a suggestive signal in AD$^{+++}$ endophenotype [*PCBD1/UNSC5B*-rs7100488; OR = 0·76 (0·69 – 0·84); p = 5·16 x $10^{-8}$]. After exploring these signals in additional datasets, we detected a nominal significance and a consistent effect for *PCBD1/UNSC5B*-rs7100488 marker only in the ADGC2 dataset [OR = 0·79 (0·61 – 1·01); p = 0·058] (Supplementary table 5).

#### Genetic exploration of GR@ACE clinical endophenotypes and enrichment analysis

For category B, with variants with stronger effects in AD mixed with vascular disease, *SORL1*-rs11218343 showed the strongest vascular enrichment [VaD$^{++}$ OR (95%) = 0·71 (0·43 - 1·16), p-value = 0·168; AD$^{+++}$ OR (95%) = 0.97 (0·77 – 1·23), p-value = 0·805], and *ADAM10*-rs593742 was the unique marker significantly associated with the VaD$^{++}$ endophenotype [VaD$^{++}$ OR (95%) = 0·80 (0·66 – 0·97), p-value = 0·02; AD$^{+++}$ OR (95%) = 0·95 (0·87 – 1·05), p-value = 0·34].

The sub-analysis of Category C allowed distinguishing two sub-categories of variants residing in this cluster. The first subset comprised variants with a stable effect across all endo-phenotypes. The second subset included variants with inverse effects between extreme clinical subgroups. Vascular processes and the regulation of nervous system development were detected in top pathways in this second subset (Supplementary Table 6)

#### eQTL for Meta-GWAS signals

To identify candidate genes and potential causal variants within novel genome-wide regions, we conducted cis-eQTL mapping. The rs2335107 marker located in the *ANKRD31* locus (chr5:74,451,443) was associated with the cortical expression of the long non-coding RNA (lncRNA) *CTD-2235C13.3* (p = 1·26 x $10^{-05}$) (appendix). This variant is located 83.4Kb from the GWAS lead SNP (rs4704171, chr5:74,368,254) and both are in complete LD ($r^2$=1). The *CTD-2235C13.3* gene is located 1·6kb from the *HMGCR* locus and its function is unknown. The *NDUFAF6*-rs4734295 marker (chr8:96,000,669), located 8.6kb from the lead meta-GWAS hit (rs10098778, chr8:95,992,020), was mapped for cortical *NDUFAF6* RNA expression (p = 1·17 x $10^{-10}$) (appendix). Finally, rs73976325 (chr17:5,123,227), located in the *SCIMP* locus 13·8kb from the meta-GWAS top signal (rs7225151, chr17:5137047), was mapped to brain cis-acting eQTL for the AC012146.1 lincRNA (p = 2·15 x $10^{-07}$) (appendix). Two additional markers, rs6502851 and rs59277121, pointed to blood eQTLs for *SCIMP* (; p = 3·89 x $10^{-08}$) and *RABEP1* (; p = 3·89 x $10^{-08}$) loci, respectively (appendix).

### References


1 Blesa R, Pujol M, Aguilar M, *et al.* Clinical validity of the 'mini-mental state' for Spanish speaking communities. *Neuropsychologia* 2001; **39**: 1150–7.

2 Blessed G, Tomlinson BE, Roth M. The association between quantitative measures of dementia and of senile change in the cerebral grey matter of elderly subjects. *Br J Psychiatry* 1968; **114**: 797–811.

3 Peña-Casanova J, Aguilar M, Bertran-Serra I, *et al.* [Normalization of cognitive and functional assessment instruments for dementia (NORMACODEM) (I): objectives, content and population]. *Neurologia* 1997; **12**: 61–8.

4 Boada M, Cejudo JC, Tàrraga L, López OL, Kaufer D. [Neuropsychiatric inventory questionnaire (NPI-Q): Spanish validation of an abridged form of the Neuropsychiatric Inventory (NPI)]. *Neurologia*; **17**: 317–23.

5 Tinetti ME, Williams TF, Mayewski R. Fall risk index for elderly patients based on number of chronic disabilities. *Am J Med* 1986; **80**: 429–34.

6 Morris JC. The Clinical Dementia Rating (CDR): current version and scoring rules. *Neurology* 1993; **43**: 2412–4.

7 Reisberg B, Ferris SH, de Leon MJ, Crook T. The Global Deterioration Scale for assessment of primary degenerative dementia. *Am J Psychiatry* 1982; **139**: 1136–9.

8 Hachinski VC, Iliff LD, Zilhka E, *et al.* Cerebral blood flow in dementia. *Arch Neurol* 1975; **32**: 632–7.

9 Alegret M, Espinosa A, Valero S, *et al.* Cut-off Scores of a Brief Neuropsychological Battery (NBACE) for Spanish Individual Adults Older than 44 Years Old. *PLoS One* 2013; **8**: e76436.

10 Román GC, Tatemichi TK, Erkinjuntti T, *et al.* Vascular dementia: diagnostic criteria for research studies. Report of the NINDS-AIREN International Workshop. *Neurology* 1993; **43**: 250–60.

11 Purcell S, Neale B, Todd-Brown K, *et al.* PLINK: a tool set for whole-genome association and population-based linkage analyses. *Am J Hum Genet* 2007; **81**: 559–75.

12 van Dam S, Cordeiro R, Craig T, van Dam J, Wood SH, de Magalhães J. GeneFriends: An online co-expression analysis tool to identify novel gene targets for aging and complex diseases. *BMC Genomics* 2012; **13**: 535.

13 Wang J, Duncan D, Shi Z, Zhang B. WEB-based GEne SeT AnaLysis Toolkit (WebGestalt): update 2013. *Nucleic Acids Res* 2013; **41**: W77–83.

14 Proitsi P, Lupton MK, Velayudhan L, *et al.* Genetic Predisposition to Increased Blood Cholesterol and Triglyceride Lipid Levels and Risk of Alzheimer Disease: A Mendelian Randomization Analysis. *PLoS Med* 2014; **11**. DOI:10.1371/journal.pmed.1001713.

15 Lourdusamy A, Newhouse S, Lunnon K, *et al.* Identification of cis-regulatory variation influencing protein abundance levels in human plasma. *Hum Mol Genet* 2012; **21**: 3719–26.

16 Naj AC, Jun G, Beecham GW, *et al.* Common variants at MS4A4/MS4A6E, CD2AP, CD33 and EPHA1 are associated with late-onset Alzheimer's disease. *Nat Genet* 2011; **43**: 436–41.

17 Filippini N, Rao A, Wetten S, *et al.* Anatomically-distinct genetic associations of APOE e{open}4 allele load with regional cortical atrophy in Alzheimer's disease. *Neuroimage* 2009; **44**: 724–8.

18 Li H, Wetten S, Li L, *et al.* Candidate single-nucleotide polymorphisms from a genomewide association study of Alzheimer disease. *Arch Neurol* 2008; **65**: 45–53.

19 Carrasquillo MM, Zou F, Pankratz VS, *et al.* Genetic variation in PCDH11X is associated with susceptibility to late-onset Alzheimer's disease. *Nat Genet* 2009; **41**: 192–8.

20 McKhann G, Drachman D, Folstein M, Katzman R, Price D, Stadlan EM. Clinical diagnosis of Alzheimer's disease: report of the NINCDS-ADRDA Work Group under the auspices of Department of Health and Human Services Task Force on Alzheimer's Disease. *Neurology* 1984; **34**: 939–44.

21 Gayán J, Galan JJ, González-Pérez A, *et al.* Genetic structure of the Spanish population. *BMC Genomics* 2010; **11**: 326.

22 Antúnez C, Boada M, González-Pérez A, *et al.* The membrane-spanning 4-domains, subfamily A (MS4A) gene cluster contains a common variant associated with Alzheimer's disease. *Genome Med* 2011; **3**: 33.

23 A. Bennett D, A. Schneider J, Arvanitakis Z, S. Wilson R. Overview and Findings from the Religious Orders Study. *Curr Alzheimer Res* 2012; **9**: 628–45.

24 Reiman EM, Webster JA, Myers AJ, *et al.* GAB2 Alleles Modify Alzheimer's Risk in APOE ε4 Carriers. *Neuron* 2007; **54**: 713–20.

25 Willer CJ, Li Y, Abecasis GR. METAL: fast and efficient meta-analysis of genomewide association scans. *Bioinformatics* 2010; **26**: 2190–1.

26 Yang J, Ferreira T, Morris AP, *et al.* Conditional and joint multiple-SNP analysis of GWAS summary statistics identifies additional variants influencing complex traits. *Nat Genet* 2012; **44**: 369–75, S1-3.

27 Bulik-Sullivan BK, Loh P-R, Finucane HK, *et al.* LD Score regression distinguishes confounding from polygenicity in genome-wide association studies. *Nat Genet* 2015; **47**: 291–5.

28 Pruim RJ, Welch RP, Sanna S, *et al.* LocusZoom: regional visualization of genome-wide association scan results. *Bioinformatics* 2010; **26**: 2336–7.

29 Machiela MJ, Chanock SJ. LDlink: a web-based application for exploring population-specific haplotype structure and linking correlated alleles of possible

functional variants: Fig. 1. *Bioinformatics* 2015; **31**: 3555–7.

Supplementary Table 1. Association results obtained from the GR@ACE dataset and GR@ACE endophenotypes, and gene categories for functional enrichment analysis.

| **Marker** | **Near Locus** | **CHR** | **BP**[a] | **Minor / Mayor** | **MAF**[b] | **VaD$^{++}$ OR (CI95%) *P*-value** | **VaD$^{+}$ OR (CI95%) *P*-value** | **Dementia OR (CI95%) *P*-value** | **AD$^{+}$ OR (CI95%) *P*-value** | **AD$^{++}$ OR (CI95%) *P*-value** | **AD$^{+++}$ OR (CI95%) *P*-value** | **Global Effect Change** |
|---|---|---|---|---|---|---|---|---|---|---|---|---|
| **Category A** | | | | | | | | | | | | |
| rs6656401 | *CR1* | 1 | 207692049 | A/G | 0·18 | 1·00<br>(1·22 – 0·82)<br>0·999 | 1·01<br>(1·15 – 0·89)<br>0·829 | 1·07<br>(1·16 – 0·98)<br>0·145 | 1·06<br>(1·16 – 0·97)<br>0·200 | 1·09<br>(1·20 – 0·99)<br>0·086 | 1·10<br>(1·22 – 0·98)<br>0·092 | 0·098 |
| rs6733839 | *BIN1* | 2 | 127892810 | T/C | 0·36 | 1·03<br>(1·21 – 0·88)<br>0·736 | 1·15<br>(1·27 – 1·04)<br>0·007 | 1·16<br>(1·24 – 1·08)<br>2·58 x $10^{-5}$ | 1·16<br>(1·25 – 1·08)<br>0·027 | 1·16<br>(1·25 – 1·07)<br>1·78 x $10^{-5}$ | 1·16<br>(1·27 – 1·07)<br>0·001 | 0·134 |
| rs190982 | *MEF2C* | 5 | 88223420 | G/A | 0·39 | 0·99<br>(1·16 – 0·85)<br>0·900 | 0·96<br>(1·06 – 0·87)<br>0·411 | 0·97<br>(1·04 – 0·91)<br>0·351 | 0·96<br>(1·02 – 0·89)<br>0·191 | 0·95<br>(1·02 – 0·88)<br>0·149 | 0·93<br>(1·02 – 0·86)<br>0·118 | 0·056 |
| rs983392 | *MS4A2* | 11 | 59923508 | G/A | 0·43 | 1·00<br>(1·17 – 0·86)<br>0·992 | 0·94<br>(1·03 – 0·85)<br>0·179 | 0·91<br>(0·96 – 0·85)<br>0·007 | 0·91<br>(0·97 – 0·85)<br>0·005 | 0·88<br>(0·94 – 0·81)<br>0·001 | 0·86<br>(0·93 – 0·79)<br>3·34 x $10^{-4}$ | 0·142 |
| rs10792832 | *PICALM* | 11 | 85867875 | A/G | 0·33 | 0·95<br>(1·12 – 0·81)<br>0·546 | 0·93<br>(1·03 – 0·84)<br>0·160 | 0·89<br>(0·97 – 0·83)<br>0·001 | 0·89<br>(0·96 – 0·83)<br>0·002 | 0·88<br>(0·95 – 0·81)<br>0·001 | 0·88<br>(0·96 – 0·81)<br>0·005 | 0·068 |
| rs2732703 | *MAPT* | 17 | 44353222 | G/T | 0·23 | 1·00<br>(1·22 – 0·80)<br>0·998 | 0·89<br>(1·01 – 0·78)<br>0·064 | 0·86<br>(0·94 – 0·79)<br>0·001 | 0·86<br>(0·93 – 0·78)<br>0·001 | 0·86<br>(0·95 – 0·78)<br>0·002 | 0·86<br>(0·97 – 0·77)<br>0·006 | 0·143 |
| rs429358 | *APOE* | 19 | 45411941 | C/T | 0·17 | 1·27<br>(1·59 – 1·02)<br>0·035 | 1·69<br>(1·93 – 1·47)<br>5·56 x $10^{-14}$ | 2·27<br>(2·50 – 2·06)<br>1·25 x $10^{-62}$ | 2·39<br>(2·64 – 2·17)<br>1·02 x $10^{-64}$ | 2·71<br>(3·01 – 2·44)<br>1·13 x $10^{-76}$ | 2·92<br>(3·27 – 2·60)<br>1·30 x $10^{-75}$ | 1·644 |
| rs3865444 | *CD33* | 19 | 51727962 | A/C | 0·29 | 0·98<br>(1·16 – 0·83)<br>0·833 | 0·93<br>(1·03 – 0·83)<br>0·173 | 0·95<br>(1·02 – 0·88)<br>0·176 | 0·94<br>(1·02 – 0·88)<br>0·134 | 0·95<br>(1·04 – 0·89)<br>0·261 | 0·92<br>(1·01 – 0·84)<br>0·067 | 0·064 |

| Marker | Near Locus | CHR | BP[a] | Mino/ Mayor Allele | MAF[b] | VaD[++] OR (CI95%) P-value | VaD[+] OR (CI95%) P-value | Dementia OR (CI95%) P-value | AD[+] OR (CI95%) P-value | AD[++] OR (CI95%) P-value | AD[+++] OR (CI95%) P-value | Global Effect Change |
|---|---|---|---|---|---|---|---|---|---|---|---|---|
| **Category B** | | | | | | | | | | | | |
| rs11218343 | *SORL1* | 11 | 121435587 | C/T | 0·03 | 0·71<br>(1·16 – 0·43)<br>0·168 | 0·79<br>(1·05 – 0·59)<br>0·109 | 0·91<br>(1·10 – 0·75)<br>0·315 | 0·90<br>(1·10 – 0·74)<br>0·310 | 0·95<br>(1·18 – 0·77)<br>0·639 | 0·97<br>(1·23 – 0·77)<br>0·805 | 0·263 |
| rs593742 | *ADAM10* | 15 | 59045774 | G/A | 0·23 | 0·80<br>(0·97 – 0·66)<br>0·022 | 0·95<br>(1·07 – 0·85)<br>0·394 | 0·94<br>(1·02 – 0·87)<br>0·149 | 0·96<br>(1·04 – 0·88)<br>0·280 | 0·95<br>(1·04 – 0·87)<br>0·267 | 0·95<br>(1·05 – 0·87)<br>0·344 | 0·150 |
| rs138190086 | *ACE* | 17 | 61538148 | A/G | 0·02 | 0·83<br>(1·00 – 0·45)<br>0·533 | 0·92<br>(1·31 – 0·64)<br>0·637 | 0·93<br>(1·18 – 0·73)<br>0·533 | 0·91<br>(1·17 – 0·71)<br>0·460 | 0·89<br>(1·18 – 0·68)<br>0·419 | 0·89<br>(1·22 – 0·65)<br>0·468 | 0·069 |
| rs11870474 | *ATP5H/KCTD2* | 17 | 73030810 | A/C | 0·03 | 1·15<br>(1·77 – 0·75)<br>0·513 | 1·43<br>(1·84 – 1·11)<br>0·006 | 1·13<br>(1·36 – 0·93)<br>0·228 | 1·13<br>(1·38 – 0·93)<br>0·213 | 1·03<br>(1·28 – 0·82)<br>0·828 | 1·08<br>(1·38 – 0·85)<br>0·510 | 0·066 |
| rs7274581 | *CASS4* | 20 | 55018260 | C/T | 0·10 | 0·92<br>(1·19 – 0·71)<br>0·526 | 0·95<br>(1·12 – 0·81)<br>0·565 | 0·96<br>(1·07 – 0·86)<br>0·435 | 0·96<br>(1·07 – 0·85)<br>0·433 | 0·97<br>(1·09 – 0·85)<br>0·588 | 0·99<br>(1·13 – 0·86)<br>0·837 | 0·066 |
| **Category C** | | | | | | | | | | | | |
| rs35349669 | *INPP5D* | 2 | 234068476 | T/C | 0·43 | 1·05<br>(1·22 – 0·90)<br>0·550 | 1·03<br>(1·14 – 0·94)<br>0·530 | 1·00<br>(1·06 – 0·93)<br>0·882 | 1·00<br>(1·07 – 0·93)<br>0·956 | 1·00<br>(1·08 – 0·93)<br>0·923 | 0·99<br>(1·07 – 0·91)<br>0·730 | 0·033 |
| rs10948363 | *CD2AP* | 6 | 47487762 | G/A | 0·27 | 1·08<br>(1·28 – 0·91)<br>0·360 | 1·10<br>(1·22 – 0·99)<br>0·084 | 1·08<br>(1·16 – 1·00)<br>0·047 | 1·08<br>(1·16 – 0·99)<br>0·063 | 1·08<br>(1·17 – 0·99)<br>0·084 | 1·08<br>(1·18 – 0·98)<br>0·125 | 0·007 |
| rs2718058 | *NME8* | 7 | 37841534 | G/A | 0·41 | 0·97<br>(1·14 – 0·83)<br>0·731 | 0·92<br>(1·02 – 0·84)<br>0·114 | 0·98<br>(1·05 – 0·91)<br>0·518 | 0·97<br>(1·04 – 0·91)<br>0·438 | 1·00<br>(1·08 – 0·93)<br>0·926 | 1·02<br>(1·11 – 0·93)<br>0·711 | 0·011 |
| rs1476679 | *ZCWPW1* | 7 | 100004446 | C/T | 0·26 | 0·98<br>(1·16 – 0·82)<br>0·774 | 0·98<br>(1·09 – 0·88)<br>0·674 | 0·94<br>(1·01 – 0·87)<br>0·103 | 0·94<br>(1·01 – 0·87)<br>0·094 | 0·91<br>(0·99 – 0·84)<br>0·038 | 0·95<br>(1·04 – 0·87)<br>0·277 | 0·025 |
| rs11771145 | *EPHA1* | 7 | 143110762 | A/G | 0·34 | 0·93<br>(1·10 – 0·79) | 0·99<br>(1·10 – 0·89) | 0·95<br>(1·01 – 0·88) | 0·95<br>(1·03 – 0·89) | 0·94<br>(1.01 0·87) | 0·92<br>(1·01 – 0·85) | 0·009 |

| Marker | Near Locus | CHR | BP[a] | Mino/ Mayor Allele | MAF[b] | VaD$^{++}$<br>OR (CI95%)<br>P-value | VaD$^{+}$<br>OR (CI95%)<br>P-value | Dementia<br>OR (CI95%)<br>P-value | AD$^{+}$<br>OR (CI95%)<br>P-value | AD$^{++}$<br>OR (CI95%)<br>P-value | AD$^{+++}$<br>OR (CI95%)<br>P-value | Global Effect Change |
|---|---|---|---|---|---|---|---|---|---|---|---|---|
| | | | | | | 0·404 | 0·837 | 0·121 | 0·202 | 0·108 | 0·071 | |
| **Category C** | | | | | | | | | | | | |
| rs28834970 | *PTK2B* | 8 | 27195121 | C/T | 0·38 | 1·06<br>(1·24 – 0·91)<br>0·461 | 1·04<br>(1·15 – 0·94)<br>0·461 | 1·04<br>(1·11 – 0·97)<br>0·283 | 1·04<br>(1·12 – 0·97)<br>0·255 | 1·04<br>(1·12 – 0·96)<br>0·317 | 1·09<br>(1·19 – 1·00)<br>0·050 | 0·028 |
| rs9331896 | *CLU* | 8 | 27467686 | C/T | 0·36 | 1·10<br>(1·28 – 0·94)<br>0·235 | 1·04<br>(1·14 – 0·94)<br>0·464 | 0·97<br>(1·04 – 0·91)<br>0·403 | 0·97<br>(1·04 – 0·90)<br>0·324 | 0·93<br>(1·01 – 0·86)<br>0·087 | 0·92<br>(0·99 – 0·84)<br>0·045 | 0·014 |
| rs10838725 | *CELF1* | 11 | 47557871 | C/T | 0·36 | 0·93<br>(1·09 – 0·79)<br>0·345 | 1·05<br>(1·16 – 0·95)<br>0·353 | 1·05<br>(1·12 – 0·98)<br>0·173 | 1·05<br>(1·13 – 0·98)<br>0·159 | 1·06<br>(1·15 – 0·98)<br>0·137 | 1·06<br>(1·16 – 0·98)<br>0·149 | 0·010 |
| rs17125944 | *FERMT2* | 14 | 53400629 | C/T | 0·07 | 0·87<br>(1·21 – 0·63)<br>0·419 | 0·96<br>(1·17 – 0·78)<br>0·665 | 1·05<br>(1·20 – 0·92)<br>0·461 | 1·05<br>(1·20 – 0·91)<br>0·515 | 1·09<br>(1·27 – 0·94)<br>0·249 | 1·19<br>(1·40 – 1·02)<br>0·031 | 0·067 |
| rs7185636 | *IQCK* | 16 | 19808163 | C/T | 0·20 | 1·25<br>(1·50 – 1·04)<br>0·018 | 1·09<br>(1·22 – 0·96)<br>0·174 | 0·97<br>(1·06 – 0·90)<br>0·547 | 0·97<br>(1·06 – 0·89)<br>0·516 | 0·93<br>(1·03 – 0·85)<br>0·148 | 0·91<br>(1·01 – 0·82)<br>0·087 | 0·159 |
| rs4147929 | *ABCA7* | 19 | 1063443 | A/G | 0·22 | 1·09<br>(1·30 – 0·90)<br>0·381 | 0·99<br>(1·11 – 0·88)<br>0·882 | 1·06<br>(1·15 – 0·98)<br>0·166 | 1·05<br>(1·14 – 0·97)<br>0·223 | 1·10<br>(1·20 – 1·01)<br>0·043 | 1·11<br>(1·23 – 1·01)<br>0·033 | 0·027 |
| rs2830500 | *ADAMTS1* | 21 | 28156856 | A/C | 0·32 | 1·05<br>(1·24 – 0·89)<br>0·537 | 1·09<br>(1·20 – 0·98)<br>0·113 | 0·99<br>(1·06 – 0·92)<br>0·810 | 0·99<br>(1·06 – 0·92)<br>0·719 | 0·95<br>(1·03 – 0·87)<br>0·198 | 0·96<br>(1·05 – 0·87)<br>0·325 | 0·009 |
| **Variants with a null effect in our dataset** | | | | | | | | | | | | |
| rs9271192 | *HLA-DRB1* | 6 | 32578530 | C/A | 0·24 | 1·01<br>(1·21 – 0·84)<br>0·897 | 0·99<br>(1·11 – 0·89)<br>0·924 | 0·99<br>(1·07 – 0·92)<br>0 ·871 | 0·99<br>(1·07 – 0·91)<br>0·793 | 0·99<br>(1·08 – 0·91)<br>0·821 | 1·00<br>(1·10 – 0·90)<br>0·942 | 0·010 |
| rs7920721 | *ECHDC3* | 10 | 11720308 | G/A | 0·40 | 1·04<br>(1·22 – 0·90)<br>0·583 | 0·99<br>(1·10 – 0·89)<br>0·983 | 1·00<br>(1·06 – 0·93)<br>0·893 | 1·00<br>(1·07 – 0·93)<br>0·914 | 0·99<br>(1·07 – 0·92)<br>0·838 | 0·99<br>(1·08 – 0·91)<br>0·891 | 0·040 |
| rs10498633 | *SLC2A4A/RIN3* | 14 | 92926952 | T/G | 0·18 | 1·02<br>(1·23 – 0·84) | 1·00<br>(1·13 – 0·88) | 0·99<br>(1·08 – 0·91) | 0·97<br>(1·06 – 0·89) | 0·97<br>(1·06 – 0·88) | 0·99<br>(1·10 – 0·89) | 0·010 |

| | | | | | | |
|---|---|---|---|---|---|---|
| | 0·873 | 0·971 | 0·851 | 0·535 | 0·490 | 0·875 |

[a] Build 37, assembly hg19. [b] Average of the entire GR@ACE cohort (n = 7,409). Global effect change = ABS(effect change Va$^{++}$ - effect change AD$^{+++}$)

Supplementary Table 2. Full description of dbGaP datasets used in the meta-analysis.

| Dataset | Cases | Controls | Quality markers for Imputation | Quality markers post-Imputation* | Genomic Inflation factor |
|---|---|---|---|---|---|
| ADGC1 | 1822 | 560 | 530,711 | 7,726,690 | 1·00 |
| ADGC2 | 599 | 239 | 531,880 | 7,720,289 | 0·98 |
| ADGC3 | 866 | 523 | 634,591 | 7,732,661 | 1·00 |
| AddNeuronMed1 | 252 | 73 | 471,757 | 7,791,484 | 0·94 |
| AddNeuronMed2 | 198 | 114 | 588,254 | 7,765,343 | 0·95 |
| ADNI1 | 258 | 73 | 620,901 | 7,704,291 | 0·94 |
| ADNI2 | 220 | 190 | 730,525 | 7,747,959 | 0·96 |
| GenADA | 785 | 764 | 374,695 | 7,589,054 | 0·99 |
| GR@ACE | 4120 | 3289 | 592,875 | 7,724,483 | 1·03 |
| MAYO | 703 | 1066 | 312,512 | 7,685,633 | 1·00 |
| NIA | 483 | 913 | 583,472 | 7,923,335 | 0·99 |
| NxC/Murcia Study | 324 | 754 | 260,399 | 7,149,673 | 1·03 |
| ROSMAP1 | 340 | 136 | 634,469 | 7,608,211 | 0·97 |
| ROSMAP2 | 214 | 58 | 730,437 | 7,559,403 | 0·93 |
| ROSMAP3 | 74 | 35 | 643,470 | 7,519,248 | 0·94 |
| TGEN | 741 | 449 | 307,631 | 7,571,653 | 1·00 |
| Meta-analysis | 11,999 | 9,236 | NA | NA | 1·10 |

*Quality markers post-Imputation: R2 > 0.30; MAF>0.01

Supplementary Table 3. eQTL analysis for novel GWAS signals associated with AD.

| **AD GWAS lead signals** | | | | **Cortical cis-eQTL** | | | | **Whole Blood cis-eQTL** | | | |
|---|---|---|---|---|---|---|---|---|---|---|---|
| **SNP** | **CHR:BP** | **Near Locus** | **LD SNPs** | **Lead Marker** | **CHR:BP** | **cis-eQTL** | **P-value** | **Lead Marker** | **CHR:BP** | **Cis-eQTL** | **P-value** |
| rs10098778 | 8: 95992020 | *TP53INP1/ NDUFAF6* | 85 | rs4734295 | 8:96,000,669 | *NDUFAF6* | $1{\cdot}17 \times 10^{-10}$ | rs4582532 | 8:95,969,007 | *TP53INP1* | $1{\cdot}17 \times 10^{-10}$ |
| rs4704171 | 5:74368254 | *ANKRD31** | 143 | rs2335107 | 5:74,451,443 | *CTD-2235C13.3* | $1{\cdot}26 \times 10^{-05}$ | NA | NA | NA | NA |
| rs7225151 | 17:5137047 | *SCIMP* | 76 | rs73976325 | 17:5,123,227 | *AC012146.1* | $2{\cdot}15 \times 10^{-07}$ | rs6502851<br>rs59277121 | 17:5,150,403<br>17:5,244,821 | *SCIMP;<br>RABEP1* | $3{\cdot}89 \times 10^{-08}$ |

**POLK* and *POC5* loci present brain non-cortical cis-e.

Supplementary Table 4. Association results for rs2732703-*KANSL1*/*MAPT* in *APOE* ε4 carriers and non-carriers across GR@ACE endophenotypes.

| | **AD**<br>**OR (CI95%);**<br>**P-value** | **AD+**<br>**OR (CI95%);**<br>**P-value** | **AD++**<br>**OR (CI95%);**<br>**P-value** | **AD+++**<br>**OR (CI95%);**<br>**P-value** |
|---|---|---|---|---|
| **APOE+** | 0·88 ( 0·81 - 0·97); 0·007 | 0·87 (0·80 - 0·96); 0·005 | 0·87 (0·78 - 0·97); 0·01 | 0·85 (0·75 - 0·96); 0·01 |
| **APOE-** | 0·88 (0·76 - 1·01); 0·07 | 0·88 (0·76 - 1·01); 0·07 | 0·87 (0·75 - 1·01); 0·07 | 0·87 (0·74 - 1·03); 0·10 |

Supplementary Table 5. Replication results for rs7100488-*PCBD1/UNSC5B* and rs117834366-*CNTNAP2* in additional datasets.

| | **rs7100488-*PCBD1/UNSC5B*** | | | | **rs117834366-*CNTNAP2*** | | | |
|---|---|---|---|---|---|---|---|---|
| | **Mayor/ Minor Allele** | **MAF** | **OR (CI95%)** | **P-value** | **Mayor/ Minor Allele** | **MAF** | **OR (CI95%)** | **P-value** |
| GR@ACE | G/A | 0.26 | 0·82 (0·76-0·89) | $4{\cdot}80 \times 10^{-07}$ | G/A | 0·01 | 2·20 (1·46-3·32) | 0·00 |
| NxC-Murcia Study | G/A | 0·27 | 1·38 (0·84-2·26) | 0·065 | NA | NA | NA (NA-NA) | NA |
| AddNeuronMed1 | G/A | 0.27 | 1·31 (0·84-2·02) | 0·232 | G/A | 0·02 | 1·76 (0·20-15·38) | 0·61 |
| AddNeuronMed2 | G/A | 0.25 | 1·05 (0·71-1·55) | 0·799 | G/A | 0·02 | 0·35 (0·05-2·52) | 0·30 |
| ADGC1 | G/A | 0.26 | 1·04 (0·89-1·22) | 0·608 | G/A | 0·02 | 0·79 (0·45-1·38) | 0·41 |
| ADGC2 | G/A | 0.26 | 0·79 (0·61-1·01) | 0·058 | G/A | 0·02 | 0·97 (0·43-2·18) | 0·94 |
| ADGC3 | G/A | 0.26 | 0·92 (0·77-1·10) | 0·384 | G/A | 0·01 | 1·27 (0·50-3·20) | 0·62 |
| ADNI1 | G/A | 0.24 | 0·99 (0·62-1·59) | 0·971 | G/A | 0·02 | 11·89 (0·44-322·73) | 0·14 |
| ADNI2 | G/A | 0.27 | 1·09 (0·78-1·51) | 0·627 | G/A | 0·02 | 0·41 (0·11-1·44) | 0·16 |
| GENADA | G/A | 0.25 | 0·94 (0·79-1·13) | 0·507 | G/A | 0·02 | 0·54 (0·24-1·25) | 0·15 |
| TGEN | G/A | 0.25 | 0·94 (0·76-1·16) | 0·582 | G/A | 0·01 | 2·22 (0·69-7·18) | 0·18 |
| MAYO | G/A | 0.25 | 1·03 (0·87-1·20) | 0·762 | G/A | 0·02 | 0·80 (0·40-1·62) | 0·54 |
| NIA | G/A | 0.26 | 0·91 (0·76-1·09) | 0·302 | G/A | 0·01 | 0·80 (0·32-1·99) | 0·63 |
| ROSMAP1 | G/A | 0·25 | 1·12 (0·81-1·56) | 0·490 | NA | NA | NA | NA |
| ROSMAP2 | G/A | 0·24 | 0·85 (0·52-1·40) | 0·524 | G/A | 0·01 | 1·97 (0·13-30·28) | 0·63 |
| ROSMAP3 | G/A | 0·20 | 0·82 (0·38-1·76) | 0·606 | NA | NA | NA | NA |
| IGAP_Stage_I | G/A | NA | 0·99 (0·95-1·02) | 0·584 | NA | NA | NA | NA |

NA: Non Available

Supplementary Table 6. Top ten biological pathways per gene cluster after sub-analysis of gene Category C.

| **GeneOntology Pathway** | **Top 10 co-regulated pathways for *Category C1*** | **P-value** |
|---|---|---|
| GO:0050865 | regulation of cell activation | $2{\cdot}22 \times 10^{-16}$ |
| GO:0002253 | activation of immune response | $3{\cdot}11 \times 10^{-15}$ |
| GO:0002764 | immune response-regulating signaling pathway | $8{\cdot}99 \times 10^{-15}$ |
| GO:0007159 | leukocyte cell-cell adhesion | $2{\cdot}08 \times 10^{-14}$ |
| GO:0009617 | response to bacterium | $3{\cdot}13 \times 10^{-12}$ |
| GO:0050900 | leukocyte migration | $6{\cdot}88 \times 10^{-12}$ |
| GO:0098542 | defense response to other organism | $2{\cdot}43 \times 10^{-11}$ |
| GO:0002274 | myeloid leukocyte activation | $3{\cdot}50 \times 10^{-9}$ |
| GO:0045785 | positive regulation of cell adhesion | $3{\cdot}96 \times 10^{-9}$ |
| GO:0060326 | cell chemotaxis | $5{\cdot}29 \times 10^{-9}$ |
| **GeneOntology Pathway** | **Top 10 co-regulated pathways for *Category C2*** | **P-value** |
| GO:0048514 | blood vessel morphogenesis | $3{\cdot}20 \times 10^{-6}$ |
| GO:0050673 | epithelial cell proliferation | $6{\cdot}54 \times 10^{-6}$ |
| GO:0051271 | negative regulation of cellular component movement | $8.74 \times 10^{-6}$ |
| GO:0040013 | negative regulation of locomotion | $1.45 \times 10^{-5}$ |
| GO:1901342 | regulation of vasculature development | $2.85 \times 10^{-5}$ |
| GO:0031346 | positive regulation of cell projection organization | $4.93 \times 10^{-5}$ |
| GO:0001655 | urogenital system development | $6.98 \times 10^{-5}$ |
| GO:0014812 | muscle cell migration | $4{\cdot}00 \times 10^{-4}$ |
| GO:0051961 | negative regulation of nervous system development | $4{\cdot}00 \times 10^{-4}$ |
| GO:0042476 | odontogenesis | $4{\cdot}00 \times 10^{-4}$ |

Supplementary Table 7. Top ten biological pathways per gene cluster after secondary gene categorization.

| GeneOntology Pathway | Top 10 co-regulated pathways for Category A | P-value |
|---|---|---|
| **Category_A[a]** | | |
| GO:0050865 | regulation of cell activation | 3.33 x 10-16 |
| GO:0032103 | positive regulation of response to external stimulus | 2.44 x 10-15 |
| GO:0002253 | activation of immune response | 3.46 x 10-14 |
| GO:0022407 | regulation of cell-cell adhesion | 7.78 x 10-14 |
| GO:0007159 | leukocyte cell-cell adhesion | 1.90 x 10-13 |
| GO:0050900 | leukocyte migration | 3.81 x 10-13 |
| GO:0045785 | positive regulation of cell adhesion | 4.27 x 10-13 |
| GO:0070661 | leukocyte proliferation | 5.15 x 10-13 |
| GO:0002764 | immune response-regulating signaling pathway | 6.29 x 10-13 |
| GO:0034341 | response to interferon-gamma | 1.32 x 10-11 |
| **Category_A[b]** | | |
| GO:1901342 | regulation of vasculature development | 2.14 x 10-7 |
| GO:0032103 | positive regulation of response to external stimulus | 4.48 x 10-7 |
| GO:0007159 | leukocyte cell-cell adhesion | 4.50 x 10-7 |
| GO:0022407 | regulation of cell-cell adhesion | 6.87 x 10-7 |
| GO:0010573 | vascular endothelial growth factor production | 1.92 x 10-6 |
| GO:0060326 | cell chemotaxis | 3.75 x 10-6 |
| GO:0050727 | regulation of inflammatory response | 1.50 x 10-5 |
| GO:0045444 | fat cell differentiation | 1.71 x 10-5 |
| GO:2001057 | reactive nitrogen species metabolic process | 4.01 x 10-5 |
| GO:0048514 | blood vessel morphogenesis | 4.82 x 10-5 |
| **Category_B** | | |
| GO:0002253 | activation of immune response | 0 |
| GO:0002764 | immune response-regulating signaling pathway | 0 |
| GO:0007159 | leukocyte cell-cell adhesion | 0 |
| GO:0050900 | leukocyte migration | 0 |
| GO:0050865 | regulation of cell activation | 1·16 x 10-16 |
| GO:0009617 | response to bacterium | 3·24 x 10-14 |
| GO:0002274 | myeloid leukocyte activation | 6·20 x 10-14 |
| GO:0060326 | cell chemotaxis | 2·23 x 10-12 |
| GO:0022407 | regulation of cell adhesion | 2·28 x 10-12 |
| GO:0031349 | positive regulation of defense | 4·72 x 10-12 |
| **Category_C** | | |
| GO:0050865 | regulation of cell activation | 3.31 x 10-9 |
| GO:0007159 | leukocyte cell-cell adhesion | 1.16 x 10-7 |
| GO:0002443 | leukocyte mediated immunity | 1.20 x 10-7 |
| GO:0098542 | defense response to other organism | 5.75 x 10-6 |
| GO:0002250 | adaptive immune response | 6.10 x 10-6 |
| GO:0032623 | interleukin-2 production | 8.03 x 10-6 |
| GO:0002764 | immune response-regulating signaling pathway | 8.58 x 10-6 |
| GO:0002683 | negative regulation of immune system process | 1.01 x 10-5 |

| | | |
|---|---|---|
| GO:0001818 | negative regulation of cytokine production | 1.53 x 10-5 |

Category A[a]: Pathway analysis conducted with top 150 co-expressed genes; Category A[b]: Pathway analysis conducted with loci co-expressing with more than 4 LOAD genes (N = 43)

Supplementary Table 8. Association results for known LOAD loci in meta-analysis of GR@ACE with dbGaP datasets and IGAP Stage I and Stages I+II.

| | | | | **dbGAP (n = 21,235)** | | | **IGAP Stage I (n = 61,571)** | | | **IGAP Stage I + II (n = 81,455)** | | |
|---|---|---|---|---|---|---|---|---|---|---|---|---|
| **Marker** | **Near Gene** | **CHR:BP** | **Major/ Minor** | **OR** | **95%CI** | **Meta *P*-value** | **OR** | **95%CI** | **Meta *P*-value** | **OR** | **95%CI** | **Meta *P*-value** |
| rs6656401 | *CR1* | 1:207692049 | G/A | 1·12 | 1·06 – 1·18 | 2·26 x $10^{-5}$ | 1·15 | 1·11 – 1·19 | 1·74 x $10^{-14}$ | 1·17 | 1·13 – 1·20 | 1·83 x $10^{-23}$ |
| rs6733839 | *BIN1* | 2:127892810 | C/T | 1·23 | 1·18 – 1·28 | 5·37 x $10^{-20}$ | 1·20 | 1·16 – 1·23 | 2·77 x $10^{-30}$ | 1·21 | 1·18 – 1·24 | 1·18 x $10^{-23}$ |
| rs35349669 | *INPP5D* | 2:234068476 | C/T | 1·03 | 0·99 – 1·08 | 0·146 | 1·05 | 1·02 – 1·09 | 6·26 x $10^{-4}$ | 1·07 | 1·04 – 1·09 | 3·45 x $10^{-7}$ |
| rs190982 | *MEF2C* | 5:88223420 | A/G | 0·93 | 0·89 – 0·97 | 0·002 | 0·93 | 0·96 – 0·90 | 3·74 x $10^{-6}$ | 0·93 | 0·96 – 0·91 | 4·00 x $10^{-8}$ |
| rs9271192 | *HLA-DRB5/ HLA-DRB1* | 6:32578530 | A/C | 1·03 | 0·98 – 1·09 | 0·214 | 1·09 | 1·13 – 1·05 | 4·51 x $10^{-7}$ | 1·10 | 1·13 – 1·07 | 1·07 x $10^{-10}$ |
| rs10948363 | *CD2AP* | 6:47487762 | A/G | 1·10 | 1·05 – 1·16 | 3·14 x $10^{-5}$ | 1·10 | 1·13 – 1·06 | 4·94 x $10^{-9}$ | 1·10 | 1·13 – 1·07 | 7·05 x $10^{-12}$ |
| rs1476679 | *ZCWPW1* | 7:100004446 | T/C | 0·92 | 0·88 – 0·96 | 4·56 x $10^{-4}$ | 0·93 | 0·96 – 0·90 | 2·07 x $10^{-6}$ | 0·92 | 0·94 – 0·89 | 1·97 x $10^{-10}$ |
| rs11771145 | *EPHA1* | 7:143110762 | G/A | 0·93 | 0·89 – 0·97 | 0·002 | 0·91 | 0·88 – 0·94 | 5·22 x $10^{-10}$ | 0·91 | 0·89 – 0·93 | 6·02 x $10^{-14}$ |
| rs2718058 | *NME8* | 7:37841534 | A/G | 0·96 | 0·92 – 1·00 | 0·040 | 0·94 | 0·97– 0·91 | 2·44 x $10^{-6}$ | 0·93 | 0·95 – 0·91 | 1·17 x $10^{-8}$ |
| rs28834970 | *PTK2B* | 8:27195121 | T/C | 1·07 | 1·02 – 1·12 | 0·002 | 1·09 | 1·12 – 1·06 | 5·81 x $10^{-9}$ | 1·10 | 1·12 – 1·07 | 1·57 x $10^{-13}$ |
| rs9331896 | *CLU* | 8:27467686 | T/C | 0·92 | 0·88 – 0·96 | 2·46 x $10^{-4}$ | 0·88 | 0·91 – 0·86 | 5·56 x $10^{-15}$ | 0·88 | 0·90 – 0·86 | 3·75 x $10^{-23}$ |
| rs11218343 | *SORL1* | 11:121435587 | T/C | 0·84 | 0·75 – 0·93 | 0·002 | 0·78 | 0·84 – 0·73 | 1·10 x $10^{-10}$ | 0·78 | 0·83 – 0·74 | 1·97 x $10^{-14}$ |
| rs10838725 | *CELF1* | 11:47557871 | T/C | 1·06 | 1·02 – 1·11 | 0·006 | 1·07 | 1·04 – 1·10 | 3·23 x $10^{-6}$ | 1·08 | 1·05 – 1·10 | 5·95 x $10^{-9}$ |
| rs983392 | *MS4A2* | 11:59923508 | A/G | 0·89 | 0·85 – 0·95 | 5·10 x $10^{-8}$ | 0·90 | 0·87 – 0·93 | 8·51 x $10^{-13}$ | 0·90 | 0·88 – 0·92 | 2·15 x $10^{-17}$ |
| rs10792832 | *PICALM* | 11:85867875 | G/A | 0·86 | 0·83 – 0·90 | 1·89 x $10^{-11}$ | 0·88 | 0·86 – 0·91 | 4·23 x $10^{-18}$ | 0·87 | 0·85 – 0·89 | 4·22 x $10^{-28}$ |
| rs17125944 | *FERMT2* | 14:53400629 | T/C | 1·10 | 1·02 – 1·19 | 0·015 | 1·12 | 1·18 – 1·06 | 1·24 x $10^{-4}$ | 1·13 | 1·18 – 1·08 | 1·10 x $10^{-8}$ |
| rs10498633 | *SLC2A4A/RIN3* | 14:92926952 | G/T | 0·92 | 0·87 – 0·96 | 7·40 x $10^{-4}$ | 0·92 | 0·88 – 0·95 | 1·24 x $10^{-6}$ | 0·92 | 0·89 – 0·95 | 3·07 x $10^{-8}$ |
| rs74615166 | *TRIP4* | 15:64725490 | T/C | 1·09 | 0·95 – 1·25 | 0·239 | 1·25 | 1·41 – 1·12 | 1·42 x $10^{-4}$ | 1·22 | 1·34 – 1·12 | 8·48 x $10^{-6}$ |
| rs2732703 | *MAPT* | 17:44353222 | T/G | 0·91 | 0·86 – 0·96 | 8·98 x $10^{-4}$ | 0·86 | 0·91 – 0·81 | 1·28 x $10^{-7}$ | 0·87 | 0·92 – 0·82 | 2·08 x $10^{-7}$ |
| rs11870474 | *ATP5H/KCTD2* | 17:73030810 | C/A | 1·13 | 1·00 – 1·28 | 0·056 | 1·17 | 1·08 – 1·27 | 2·81 x $10^{-4}$ | 1·16 | 1·07 – 1·25 | 1·11 x $10^{-4}$ |
| rs4147929 | *ABCA7* | 19:1063443 | G/A | 1·08 | 1·02 – 1·15 | 0·008 | 1·12 | 1·08 – 1·17 | 2·83 x $10^{-9}$ | 1·14 | 1·10 – 1·17 | 2·44 x $10^{-15}$ |
| rs429358 | *APOE* | 19:45411941 | T/C | 3·21 | 3·02 – 3·41 | 1·88 x $10^{-303}$ | 3·41 | 3·57 – 3·25 | 0 | NA | NA | NA |
| rs3865444 | *CD33* | 19:51727962 | C/A | 0·90 | 0·86 – 0·94 | 6·75 x $10^{-6}$ | 0·92 | 0·89 – 0·95 | 3·61 x $10^{-8}$ | 0·94 | 0·91 – 0·96 | 1·13 x $10^{-6}$ |
| rs7274581 | *CASS4* | 20:55018260 | T/C | 0·87 | 0·81 – 0·93 | 1·02 x $10^{-4}$ | 0·89 | 0·93 – 0·84 | 3·92 x $10^{-6-}$ | 0·89 | 0·93 – 0·85 | 5·17 x $10^{-8}$ |
